# Expressed therapeutic protein yields are predicted by transiently transfected mammalian cell population

**DOI:** 10.1101/2022.03.15.484372

**Authors:** Ly Porosk, Jekaterina Nebogatova, Heleri Heike Härk, Birgit Vunk, Piret Arukuusk, Urve Toots, Mart Ustav, Ülo Langel, Kaido Kurrikoff

## Abstract

Therapeutic proteins are currently at the hotspot of innovation in the pharmaceutical medicine. However, their industrial production is technically challenging and improved methods for transcriptional manipulation of mammalian industrial cell cultures are needed. In this work we show that some of the most frequently used lab scale transfection efficacy assays fail to predict performance in the protein production settings. We compare the efficacies of a number of transfection reagents using adherent and suspension mammalian cell cultures and assessment based on several assays that utilize reporter protein quantitation, transfected cell population and post-transfection viability of cells. We validate reporter assays for assessing transfection methods in the lab that predict protein production in industrial settings. We also demonstrate that cell penetrating peptide-based transfection achieve significantly higher protein yields compared to PEI and lipoplex methods in both CHO and HEK293 producer cell lines. Availability of fast lab scale screening methods allows future development of improved transfection methods for protein production. One such potentially effective transient transfection method is the CPP-based approach presented currently.

## 1. Introduction

Therapeutic proteins represent a huge and growing field in the pharmaceutical medicine: half of the global top 10 revenue drugs sold in 2021 were either humanized monoclonal antibodies (mAb) or other protein-based biologicals [1]. Therapeutic intervention using mAb has been very successful, especially in cancer and inflammatory diseases. mAb allow treatment with improved efficiencies and lower side effects at a magnitude that has not been achievable with chemically synthesized small-molecule drugs [2].

With increasing demand for recombinant proteins, there is growing need for rapid methods to produce the protein of interest. The industrial production of protein drugs is considerably more challenging than production of chemically synthesized compounds and thus improvement of the current cell culture manipulation methods has huge importance [3]. Protein drugs are produced in cultivated mammalian cells in order to ensure their proper folding, assembly and post-translational modifications. One of the challenges that determine protein production yield is the process of transfection where plasmid DNA (pDNA) expression vector is introduced into the cells in order to use the cell’s protein biosynthesis machinery.

Transient transfection was introduced for the mass use (with scale-up compatibility) in 1995 with polyethylenimine (PEI) method [4]. Since then, transient transfection is the first choice for rapidly obtaining recombinant proteins in industrial settings without the need to spend time and resources for stable cell lines[5]. The technology has leaped forward with the use of cell engineering and culturing advancements-Protein yields that used to be in the range of mg/L 20 years ago, have now increased to an impressive 1–2 g/L in CHO and HEK293 producer cell lines [3].

Nevertheless, efficient transfection and subsequent protein expression cannot be achieved through a universal method and the outcome is influenced by many aspects, such as used cell lines,culturing conditions etc[3]. There are literally hundreds of different chemical transfection reagents available to choose from, but these have been generally developed for the laboratory scale. In this work we aim to assess how well the lab scale efficacy estimation holds up when scaling up and how well we can predict protein expression success in industrial settings. We subsequently propose lab assays with better predictive power for efficient protein production.

## 2. Results

### 2.1. Classically used transfection efficacy assays fail to predict protein production in industrial settings

The most frequently used transient transfection methods are polyplex (PEI) and lipoplex-based approaches [5] and typical way of assessing the transfection efficacy is by quantitation of the expressed reporter. Frequently, total luciferase amount is measured from the cell lysate in an adherent cell culture at 24…48h post-transfection. Luciferase reporter assay is sensitive, fast, requires simple equipment and is widely available, and since the desired outcome in the industry is maximizing the yields of protein-of-interest, it seems reasonable to focus on quantitation of total reporter protein.

Here we compared pDNA transfection efficacy of a small set of well-known lipoplex, polyplex, as well as cell-penetrating peptide (CPP)-based transfection methods (Table 1) in CHO cells, which is the main industrial producer cell line [3,6]. Since each transfection reagent has its own instructions for use, in many cases leaving room for optimization, assay conditions were selected by optimizing the reagent amount (Fig S 1a), N/P ratio (Fig S 1b), charge ratio (Fig S 1c), media change and pDNA dose (Fig S 2a-c) to maximize the transfection output. As a result, we observed generally high protein expression with all the methods, although lipoplex and polyplex methods were the most effective ones (Figure1a). CPP-mediated transfection, albeit efficient, did not quite reach the levels shown by jetOptimus, Lipofectamine nor PEI-Max (statistical analysis in Supplementary Table 2).

**Figure 1.**
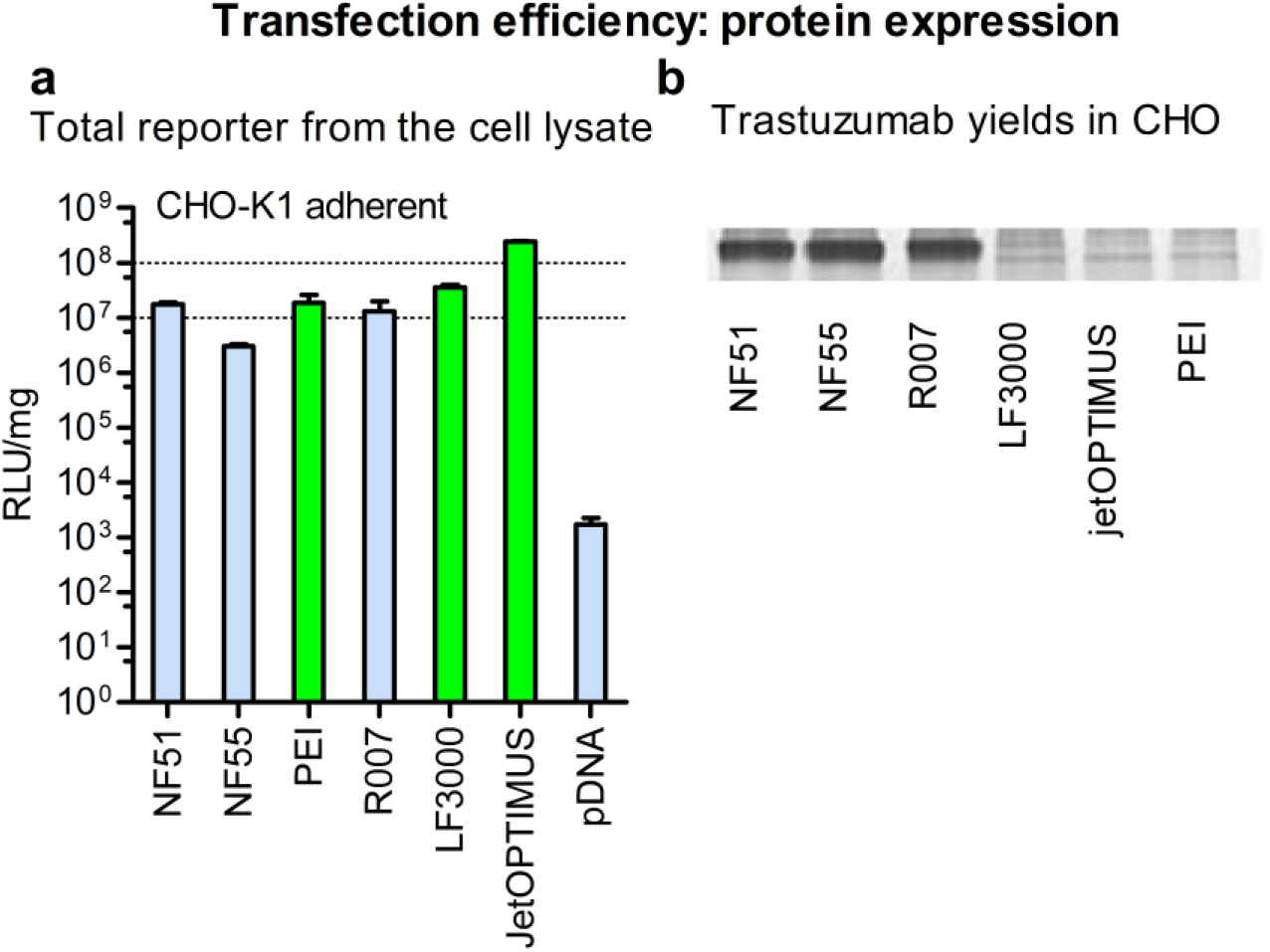
Total expressed reporter protein levels in adherent cells do not correlate with therapeutic protein production efficacy. **a)** Transfection efficacy assessed by firefly luminescence values (RLU) from cell lysate 24h post-transfection of CHO-K1 adherent cells. RLU normalized to total protein. The top performers are highlighted with green. **b**) Production of Trastuzumab mAb in CHO 1H7 suspension cell culture in serum free defined media 7 days after transient transfection of pDNA. Assessed by SDS-PAAG 10%, coomassie staining.

**Table 1..** Transfection methods and their classifications

| Name* | Abbreviation | Classification (if known) |
| --- | --- | --- |
| NickFect51 [8] | NF51 | CPP |
| NickFect71 [11] | NF71 | CPP |
| PepFect14 [12] | PF14 | CPP |
| NickFect55 [9] | NF55 | CPP |
| PEI-MAX | PEI | polyplex |
| Reagent007 | R007 | CPP |
| Lipofectamine 3000 | LF3000 | lipoplex |
| Turbofect |  | lipoplex |
| Xfect |  | polyplex |
| JetOPTIMUS |  | Cationic transfection reagent |
| TransIT X2 |  | Non-liposomal polymer |

We were next interested in how well these performances translated into the “real” protein expression settings. We therefore undertook the production of an example protein, therapeutic mAb Trastuzumab by using a QMCF protein expression technology [7] in CHO suspension cells over a 1-week production cycle. The QMCF includes EBNA and PyLT elements in the producer cells and plasmid expression vectors with the aim of enhancing the retention and replication of the protein expression vector in the cells and allowing significant extension of the “transient” period in the protein production. Other protein expression methods with similar mechanisms are also available [3]. We expected to find that all the transfection methods were able to produce reasonable yields of mAb, especially the top performers from the Figure1a, since that was indicated by high reporter protein expression from the same cell line at short time frame.

Surprisingly, however, we observed drastic differences from the expected: the CPP methods NF55[8], NF51 [9] that we previously described, as well as the Reagent007 significantly outperformed both the lipoplex and polyplex approaches (Figure1b). More importantly, we concluded that the fast and easy luciferase reporter lab assay fails to predict industrial protein production performance. Considering that therapeutic protein production is technically challenging and the final yields are defined by the success of cell culture transfection step, the whole field would benefit from being able to predict large scale performance after weeks of production in cell factories [10]. Availability of fast lab scale screening methods would allow development of improved transfection methods for protein production.

In order to explore the transfection efficacies and possibilities for protein production further, we expanded our selection to include more transfection methods, such as our previously described CPP transfection methods PF14 [12] and NF71 [11], but also other known examples of polymers and liposomes. Furthermore, to explore if the above described discrepancy is specific to CHO, we also compared luciferase expression performances in another cell line. HEK293 is well-established producer cell line because of its ease of use for both cell growth and transfection, as well as its excellent protein yields. It is frequently utilized as the expression system for the production of recombinant proteins and viruses for gene therapies [5]. Again, to be able to compare different reagents, optimal conditions were determined to maximize the transfection result (Fig S 1a-c, Fig S 2a-c). Transfection efficacies with this larger set were in line with what we observed in Figure1a: the top performers in both settings were liposomal and polymeric approaches with statistically significantly higher values (Figure2a, blue bars, top 3 are highlighted in green, statistical analysis in Supplementary table 2). As for the HEK293 cell line, the differences between the methods were smaller, but liposomal and PEI-based approaches were still the top performers (Figure2a, dark bars).

**Figure 2.**
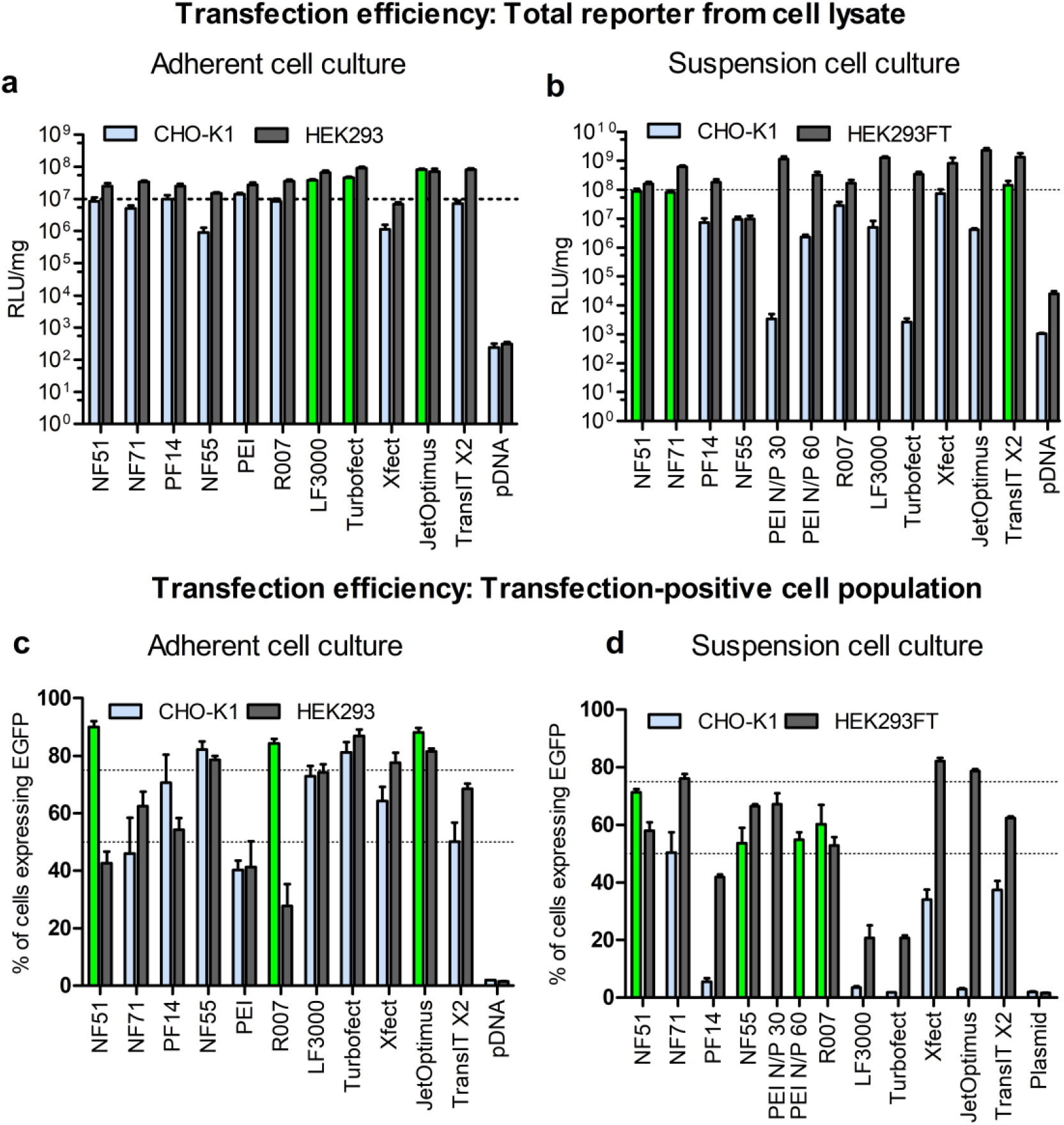
High expression of reporter protein can be achieved with most transfection methods. **a**) Total reporter protein levels from cell lysate of adherent cells. Measurement 24h post-transfection. Transfected with 0.2 μg of firefly luciferase encoding pLuc per 96-well plate well. Transfection in serum free media and 4h post-transfection media was changed to serum-containing media. CPP/pDNA complexes formed at CR3, PEI N/P 20 **b)** Total reporter protein levels from cell lysate of suspension cells transfected with 0.1 μg of pDNA per 96-well plate well. CPP/pDNA complexes formed at CR2. Measurement 48h post-transfection in serum-free media. The results are shown as relative luminescence units (RLU) normalized to total protein (mg) from lysate. **c**) Adherent cells transfected with 0.5 μg of green fluorescent protein encoding plasmid (pGFP) per 24-well plate well. Transfection in serum free media and 4 h post-transfection media was changed to serum containing mediaCPP/pDNA complexes were formed at CR3, PEI N/P 20. **d)** suspension cells transfected with 0.75 μg of pGFP per 24 well-plate well. Transfection in serum free media. CPP/pDNA complexes formed at CR 2. 4 h post-transfection 500 μl of fresh media was added to each well. For PEI N/P20 was used in case of HEK293FT cells and N/P60 in case of CHO-K1 cells.48 h post-transfection cells were analyzed by flow cytometry.

### 2.2. Using suspension culture improves the predictability for protein production

An important difference in the industrial cell factories is the use of mammalian suspension cultures, where much higher cell densities are achieved than it is possible in adherent cell cultures. Hence, we next measured transfection efficacies in suspension culture by using the same total luciferase quantitation assay. Transfection time and volume, presence of serum in the media, transfection reagent dose, pDNA dose, and N/P ratio were optimized to maximize the expressed protein levels (Fig S 3a-b, Fig S 4a-d). The general observation of the resultant efficacies was that the distinction between high and low performers was more pronounced than in adherent conditions (Figure2a vs b). The most notable was that the top performers in CHO suspension culture were completely different from what we saw in adherent conditions, and now included 2 peptide-based methods (NF51 and NF71) and one polyplex (TransIT) method (Figure2b, highlighted in green). Importantly, when comparing the high and low performers in suspension culture to the initial set of methods included in the mAb production (Figure1b), the predictive value of suspension culture transfection is higher than what is observed in adherent cell culture. One of the efficient protein producers—NF51—was among the top 3 transfection methods, and conversely, none of the nonperformers in mAb production were among the best transfection methods in CHO suspension culture (Figure2b vs Figure1b, statistical analysis in Supplementary Table 2).

### 2.3. Transfected cell population is an important efficacy predictor

Although the amount of total expressed protein (as shown in Figure2) may be intuitively correct way to estimate transfection performance, long-term protein production is also dependent on another important aspect—successful division of transfected cells. Each transfection-positive cell clone gives rise to approximately 100 to 500 protein-producing offspring, depending on the cell line and cultivation conditions [13]. For example, a cell that is successfully transfected (estimated at early time point, such as in Figure2) but is unable to undergo cell division and produce daughter cells (which will only be apparent after several days post-transfection, such as in Figure1b) is ultimately not contributing to expressed protein yields.

To assess the effect of successful transfection of many vs few cells, we followed two separate lines of reasoning. To get an idea about the cells that contribute towards successful transfection, we measured the transfection-positive cell population. On the other side, to observe the cells that fail to contribute to transfection, we studied the number of live/dead cells post transfection.

First, we measured and compared the transfection-positive cell population by switching to the EGFP reporter and counting fluorescent-positive cells with the cell counter after transfection with pGFP. We validated that both cell lines retained the fluorescent signal from the expressed plasmid beyond 5 days (Fig S6a and b) and 48h post-transfection was chosen for this assay (Fig S 6c). We completed this assessment first in adherent cells and observed that indeed, the predictability for protein yields had further increased: now 2 out of the top 3 protein producers in CHO—NF51 and Reagent007—were among the top transfectors (from Figure1b), and the third—NF55 was almost at the level of the best (Figure3a, statistical analysis in Supplementary Table 2). However, there is still room for improvement, because some of the nonperforming protein producers were also flagged among the efficient transfection methods (such as jetOptimus in CHO in Figure3a).

**Figure 3.**
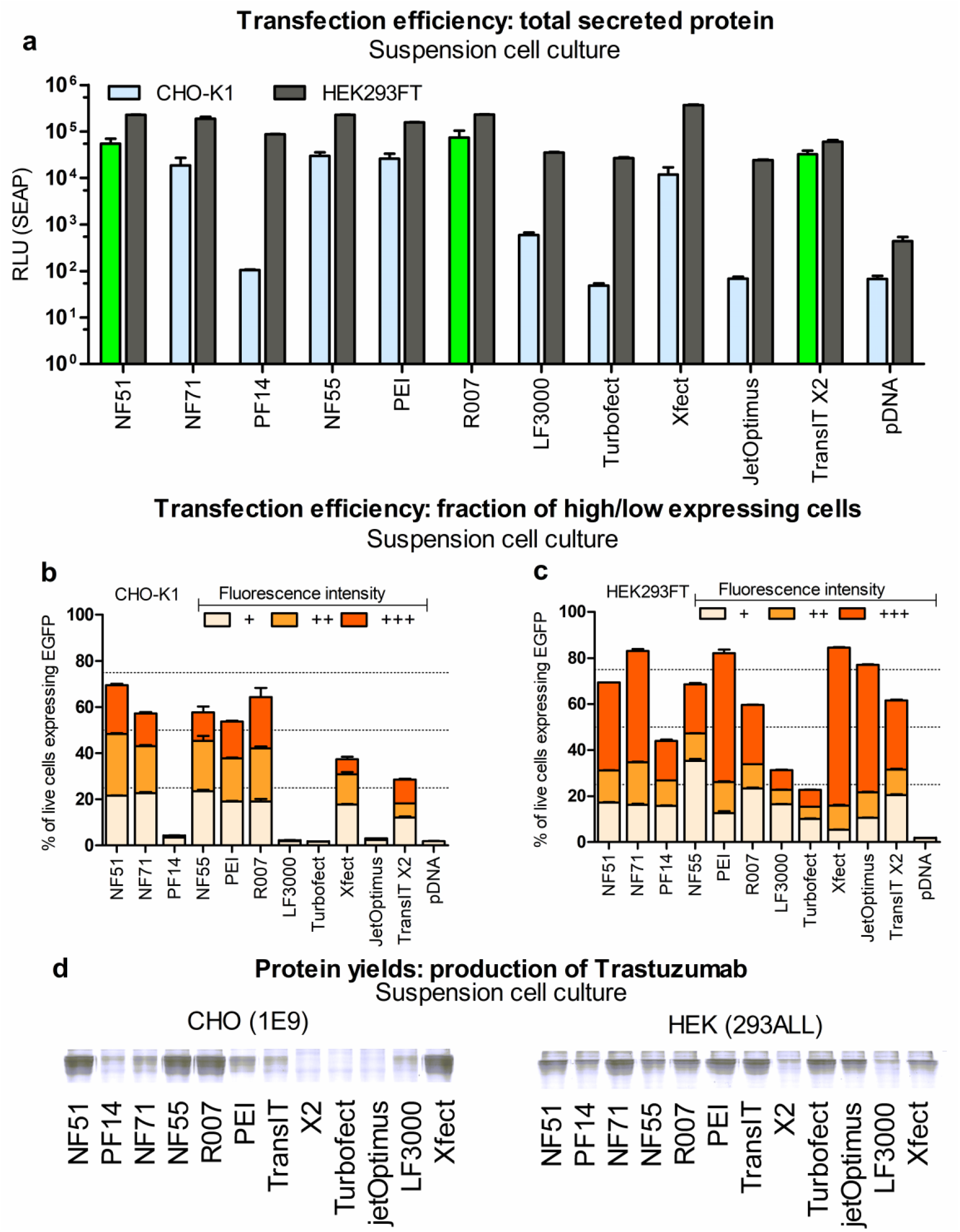
Transfection efficacy in suspension cell culture indicates possible yields of expressed Trastuzumab mAb. a-c) CHO-K1 and/or HEK293FT cells transfected in serum free media with 0.75 μg of pDNA. 4 h post-transfection fresh media was added. Analysis was performed 48 h post-transfection. CPP/pDNA complexes were formed at CR2. PEI N/P 20 (HEK293FT) or N/P60 (CHO-K1). **a)** Cells transfected with pSEAP. 48 h post-transfection luminescence from SEAP was detected from media. **b**) CHO-K1 and c) HEK293FT cells transfected with 0.75 μg of pGFP. The fluorescent positive cell population was detected 48 h post-transfection from cell population. The fluorescence intensities of fluorescent positive cells were divided into low signal (+), medium signal (++) and cells with strong signal (+++) fractions. Suspension **d)** Trastuzumab mAB production in suspension cell culture. 2 μg of pLic2.1 pDNA per 6-well plate well was used. For CHO 1E9 cells analysis was performed 8 days post-transfection. For HEK293ALL analysis was performed 5 days post-transfection. Assessed by SDS-PAAG 10%, coomassie staining.

Our next question was: does transfection-positive cell count improve protein production prediction if we use assay in suspension culture, as we observed in case of total luciferase quantitation? Transfection-positive cell count profiles are presented in Figure3b. In order to fully compare and analyze potential correlations with the protein production, we included all the transfection methods listed in Figure 2 and Figure 3 side by side in the production of mAb in another independent experiment. We did this both in CHO as well as HEK293 producer cells over a 1-week production cycle (Figure3c–d).

The CHO production profile did not reveal further surprises besides the initial observation that we already discussed above in Figure1b, that the polymeric and liposomal methods were inferior to the peptide-mediated approaches (Figure3c). It is noteworthy that only some of the CPPs were apparently effective for mAb production, for example, PF14, being an excellent transfection method that has been repeatedly demonstrated previously [14,15], performed poorly through the whole current set: transfection of the adherent (Figure3a) and suspension culture (Figure3b), protein production in both CHO (Figure3c) and HEK293 (Figure3d). It should be noted that this poor performance is not in a disagreement with the previous reports, as PF14 was specifically developed for the transfection in serum-containing media [12] and with *in vivo* gene therapeutic applications in mind [16-18], whereas in the current report, performances are strictly compared in serum-free, defined media that is routinely used in the applications of protein production in cell factories [3]. It is probable that the other non-performers shown in Figure 3 were also developed for general lab-use in mind. The message here is that in order to improve the application of mammalian transfection in cell factories, the reagents should be specifically developed for that goal and using proper assays to estimate efficacies may help accomplish that goal.

The performances in CHO cell line assessed by measuring positive cell population with flow cytometry (Figure3b) seem to successfully flag both performers (highlighted in green) and non-performers in mAb production: NF51, NF55, Reagent007 and PEI-Max are among the best transfectors as well as the protein producers (although PEI-Max can not be considered on par with the CPP methods), whereas the rest of the methods lack both in transfection and mAb production (statistical analysis in Supplementary Table 2).

In HEK293, the protein is produced most effectively with the CPP NF71, followed by polyplex methods PEI-Max, TransIT, and jetOptimus (Figure3d). However, neither of the reporter protein assays (total protein or transfection-positive cell population) and neither of the culturing conditions (adherent or suspension) were able to predict performance in mAb production, so further investigation is needed for this cell line.

### 2.4. Expressed protein yields are predicted by total secreted protein reporter

In order to analyze if specific subpopulations of cells with high or low expressed protein levels determine the outcome of the ultimate protein yields, we took a closer look at the GFP+ cells. We defined transfection-positive cell subpopulations into 3 subcategories: cells with low, medium, and high expressed protein levels (Fig S 7, Fig S 8, Fig 3b-c). However, none of these subfractions allowed making more useful predictions, because the subpopulation proportions correlated mostly only with the total GFP+ cells (Supplementary Table 3).

Considering that we are interested in effective producing of large mammalian proteins, including mAb that are processed in ER and Golgi, and are secreted, we included yet another reporter protein that is also critically dependent on the ER and Golgi processing and is secreted out of the cell—the secreted alkaline phosphatase (SEAP). The profiles of pSEAP transfection are shown in Figure3c (statistical analysis in Supplementary Table 2). Importantly, we observed that the SEAP expression in CHO suspension culture correlated strongly with the mAb protein yields (r=0.88, p < .05). In HEK293 suspension culture, SEAP expression correlated with the number of GFP+ cells, but unlike in CHO, neither of these efficacy assays correlated with mAb production (Supplementary Table 3).

### 2.5. High viability of cells does not correlate with high protein expression

It is obvious that successful expression of proteins is dependent on the number of viable cells. In order to characterize the cells that fail to contribute towards transfection, we analyzed the number of live and dead cells. Although we also considered the BrdU (Fig S 9a) and MTS assays (Fig S 9b), the Live/Dead assay has the advantage that it is based on the combination of Calcein AM and Propidium iodide (PI) staining and reflects both live cell and cells with compromised cell membrane in a cell population (Fig S 10, Fig S 11, Fig S 12). Our intention was to analyze if indeed the proportion of (living) cells correlated with expressed protein amount and if we could observe higher number of cells in the groups that perform well in protein production.

In the CHO adherent cells, we observed that some transfection methods were accompanied by apparent reduction of viable cells. Controversially, the most “toxic” methods were concurrently very efficient transfection mediators in adherent culture: the Pearson coefficient between adherent CHO total luc expression and number of live cells was r=-0.85 (p < .05). We exemplified two different cases from this curious correlation and visualized these with the CLSM: jetOptimus as the No1 in total luc transfection, but also the one with the least number of live cells (Supplementary Table 3). On the other hand, among the current selection of methods, NF55 is the least performing in total luc expression in adherent CHO, but among the group with the highest number of viable cells (Figure4c). On one hand, we can conclude that reduced cell viability post-transfection may not mean much in adherent conditions, i.e. it is not necessarily a negative occurrence by itself. However, in the context of protein production in defined media, total luciferase expression in adherent cell culture definitely is not conjoined with the protein yields, if anything, the correlation is negative (r=-0.59, but statistically not significant).

**Figure 4.**
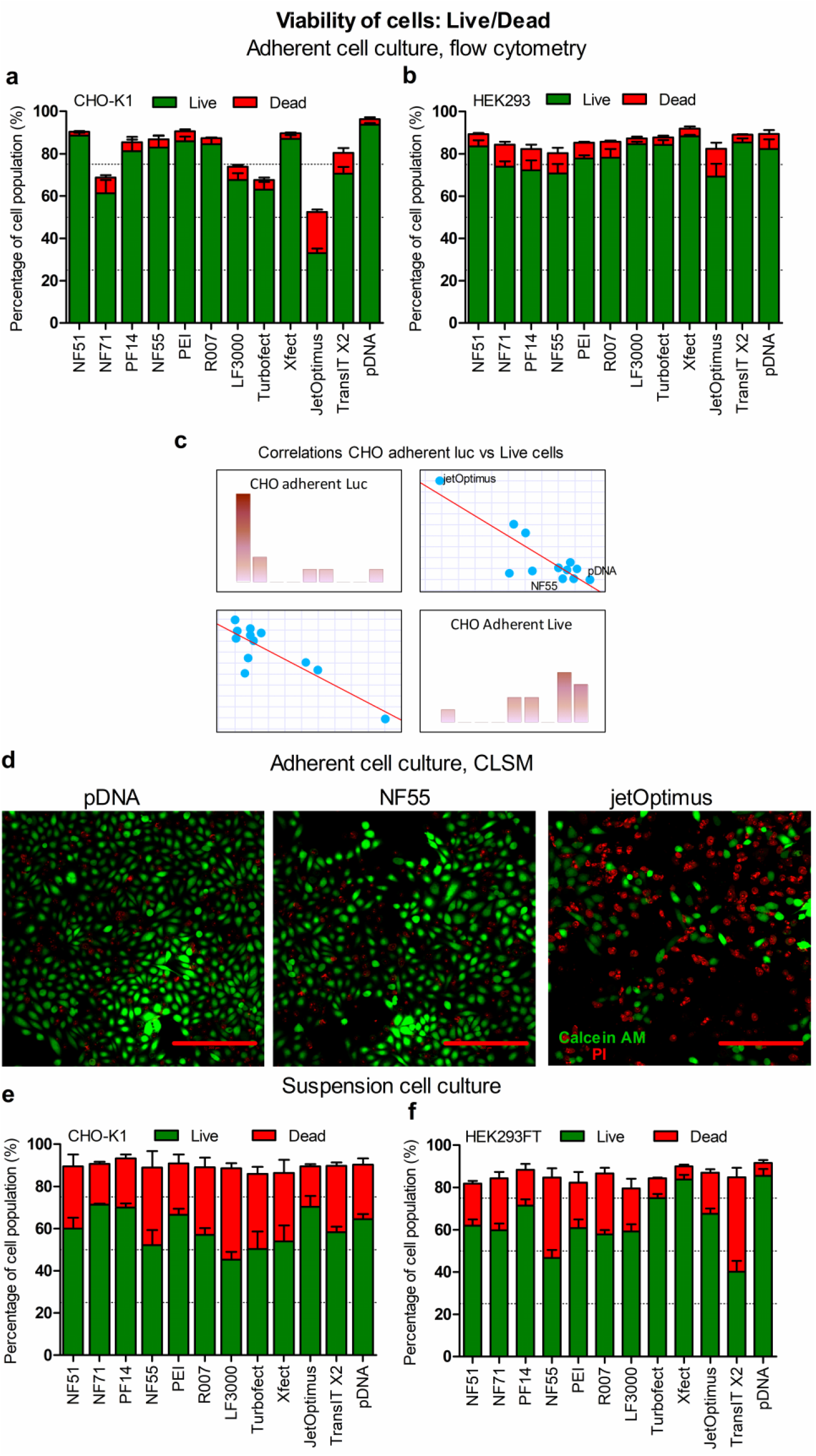
Cell viability post-transfection assessed by Live/Dead assay. Adherent **a**) CHO-K1 cells and **b**) HEK293 cells transfected with 0.5 μg of pLuc per 24-well plate well, in serum free media. CPP/pDNA complexes formed at CR3, PEI N/P20. 4 h post-transfection media was replaced with serum containing media24 h post-transfection Calcein AM (Live) and PI (Dead) were used to detect Live and Dead cells from cell population. c)Correlation graphs depicting correlations between luciferase expression levels against viability of cells. Adherent cell experiments. d) Representative confocal images of cells treated with pDNA, cells transfected using jetOPTIMUS (jetOptimus) or NF55 (NF55) Red bar corresponds to 200 μm. Analysis 24 h post-transfection. Suspension **e**) CHO K1 and **f**) HEK293FT cells transfected with 0.75 μg of pSEAP. CPP/pDNA complexes formed at CR2, PEI N/P20 (HEK293FT) or N/P60 (CHO-K1). 4 h post-transfection equal volume of fresh media was added. 48 h post-transfection cells were collected, andCalcein AM and PI were added to detect Live and Dead cells from cell population.

**Figure 5.**
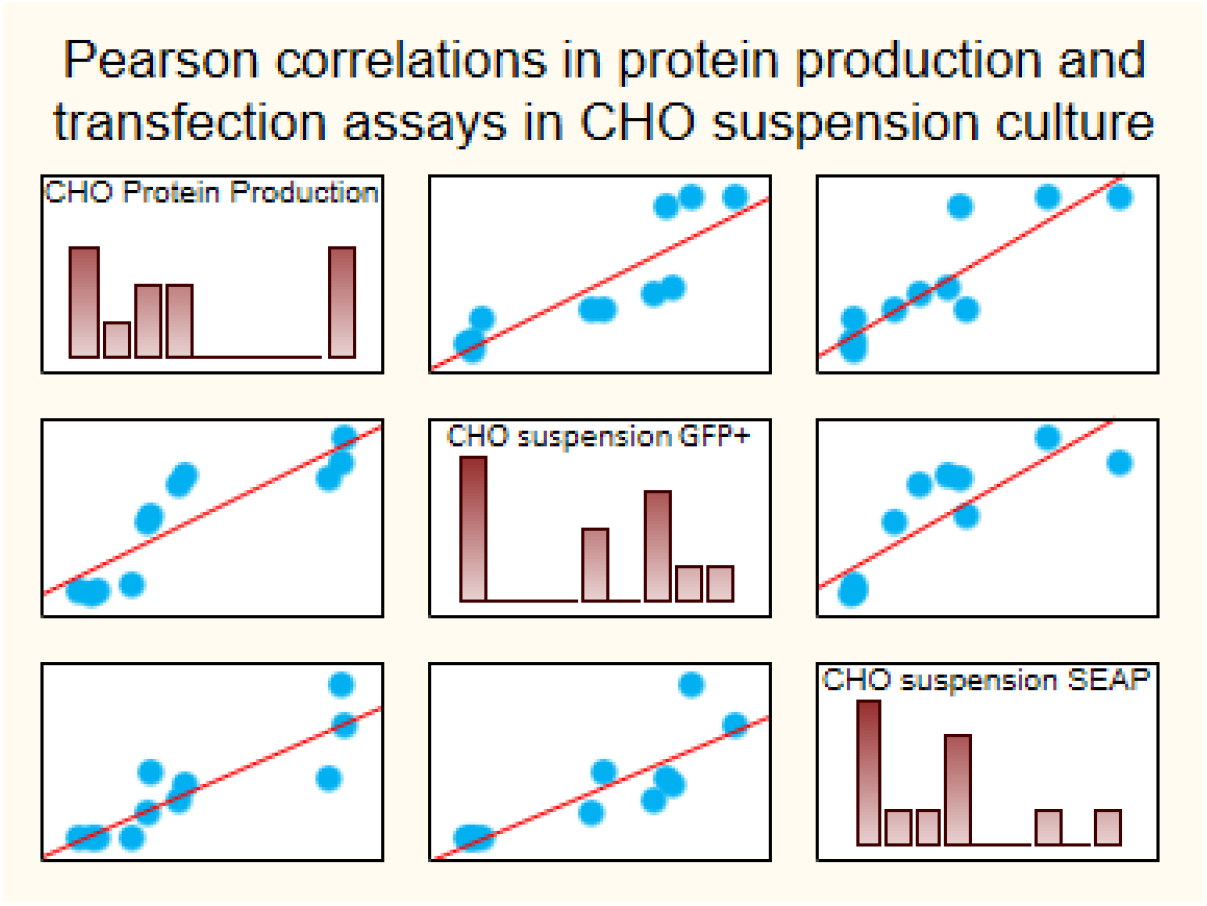
The protein production yields correlate with short term assays of secreted reporter protein and transfection+ population in CHO suspension culture. A scatterplot that highlights the correlations between the protein production yields and the assays of transfection-positive cell population (r=0.86, p < .05) and SEAP protein secretion (r=0.88, p < .05) in CHO suspension cells. The full correlation matrix with the Pearson coefficients is presented in **Supplementary Table 3**.

## 3. Materials and Methods

### 3.1 Transfection reagents

Cell-penetrating peptides (CPP) NickFect51 (NF51), NickFect55 (NF55), NickFect71 (NF71) and PepFect14 (PF14) (sequences shown in Supplementary Table 1) were developed for *in vitro* and *in vivo* nucleic acid delivery, and represent four different design strategies for PF and NF CPP lines. NF51 was chosen as it has been shown to efficiently deliver different nucleic acid cargoes into a variety of cell lines, including hard to transfect cells [7], both in serum free and serum containing conditions. Its further development, with *in vivo* delivery in mind was NF55, with an advantage in serum containing conditions and it has been used for pDNA delivery *in vivo* mouse models [8]. It was chosen because of its shown efficacy as plasmid delivery vector. In order to gain triggered release of delivered cargo, further carriers were developed, and one of them was NF71 [10].NF71 was chosen as it has been shown to efficiently deliver nucleic acids into cells, including hard-to-transfect cells. PF14 is a CPP rich in ornithines and leucines, with relatively high positive net charge. Ithas been used to efficiently deliver pDNA *in vitro* and *in vivo* [11].

CPPs were synthesized on an automated peptide synthesizer (Biotage Initiator+ Alstra) using the fluorenylmethyloxycarbonyl (Fmoc) solid phase peptide synthesis strategy with Rink-amide ChemMatrix resin (0.41 mmol g−1 loading) to obtain C-terminally amidated peptides. The fatty acid was coupled manually to the N-terminus of the peptide overnight, at room temperature with 5 eq. fatty acid. For the synthesis of the NF51, NF71 and NF55 the Boc-L-Orn(Fmoc)-OH (Iris Biotech, Germany) was used to continue the synthesis from the side-chain amino group. The reaction was carried out in DMF using HOBT/HBTU for manual or DIC/Oxyma for machine synthesis as coupling reagents, with DIEA as an activator base. Cleavage was performed with trifluoroacetic acid, 2.5% triisopropylsilane and 2.5% water for 2 h at room temperature. Peptides were purified by reversed-phase high-performance liquid chromatography on a C4 column (Phenomenex Jupiter C4, 5 μm, 300 Å, 250 × 10 mm) using a gradient of acetonitrile/water containing 0.1% TFA. The molecular weight of the peptides was analyzed by matrix-assisted laser desorption-ionization/time of flight mass spectrometry (Brucker Microflex LT/SH, USA). The concentration of the peptides was determined based on dilutions of accurately weighed substances and absorption of tyrosine, where applicable.

Commercially available transfection reagents included in this work were linear polyethyleneimine based Polyethylenimine HCl MAX (PEI), MW 40,000 from Polysciences. Peptide-based transfection reagent Reagent007 from Icosagen suitable for suspension cells. From Thermo Fisher Scientific two reagents, Lipofectamine™3000 (LF3000) Transfection Reagent as a lipid nanoparticle based technology, and Turbofect as a lipid-based cationic polymer transfection reagent were chosen. LF3000 and Turbofect are marked as suitable for nucleic acid transfection in a variety of cell lines. The LF3000 transfection system contains transfection reagent and enhancer for pDNA delivery. Xfect from Takara Bio is characterized as a biodegradable transfection polymer suitable for both CHO and HEK, and adherent or suspension cells. From Polyplus we chose jetOPTIMUS which is characterized as a cationic transfection reagent suitable for transient and stable gene expression in mammalian cell culture. It should be noted, that as a recent addition to the manufacturer’s portfolio FectoPRO was introduced specifically for protein production. From Mirus Bio we chose TransIT X2 characterised as a non-liposomal polymeric transfection system.

### 3.2 Cell culture maintenance

Adherent CHO-K1 (ECCC, CHO 85050302) and HEK293 (Obtained from Prof. Andres Merits) cells were grown in Dulbecco’s Modified Eagle’s Medium (DMEM), supplemented 0.1 mM non-essential amino acids, 1.0 mM sodium pyruvate, 100 U/ml penicillin, 100 mg/ml streptomycin. For complete media 10% (final) fetal bovine serum (FBS) was added. For transfection, serum free media was used. Cells were passaged regularly when the confluence of cells reached 80-90%. Cells were maintained in a humidified incubator at 37°C, 5.0 % CO_2_. Suspension CHO-K1 cells were adapted to suspension culture, and HEK293FT cells (Obtained from Dr. Alla Piirsoo, Thermo Fisher Scientific; catalog number: R70007) cells were grown as suspension cell culture in Xell HEK TF media, supplemented with 100 u/ml penicillin, 100 mg/ml streptomycin and 6 mM Glutamax. Cell viability was assessed daily and cell density was reduced regularly. Cells were maintained in a humidified incubator at 37°C, 8.0 % CO_2_. Suspension CHO 1E9 and 293ALL cells were grown as suspension culture. CHO 1E9 cells were grown in Xell CHO TF media. 293ALL cells were grown in BalanCD HEK293 or Xell HEK TF media. Media was supplemented with 4 mM GlutaMAX. For cell counting CytoSMART cell counter accompanied by 0.4% trypan blue staining prior measurement was used. The multi-well plates with cell cultures used for experiments were incubated in a humidified incubator at 37°C, 5.0 % CO_2_ for adherent cell cultures and at 37°C, 8.0 % CO_2_ for suspension cell cultures. For Trastuzumab monoclonal antibody expression cell cultures were incubated in humidified incubator at 37°C, 8.0 % CO_2_ on an orbital shaker platform.

### 3.3 Transfection

Four different reporter plasmids were used throughout this work. Firefly luciferase encoding plasmid (pMC.BESPX-GLucFLuc2, referred as pLuc), pEGFP-C1 plasmid (pGFP) expressing green fluorescent protein, and pCMV_SEAP (pSEAP) expressing secreted alkaline phosphatase. SEAP expressing plasmid pSEAP was generated by substitution of eGFP coding sequence in pEGFP-C1 with sequence coding for SEAP in pLIVE® In Vivo Expression/Reporter Vectors (Mirus MIR 5620). SEAP integration into pSEAP plasmid sequence was checked by sequencing. For protein production Trastuzumab monoclonal antibody expressing pLic2.1 plasmid was used.

Plasmid doses and transfection volumes are defined under specific experiments. For CPP/pDNA complex formation the diluted pDNA and peptide were mixed in milliQ water. Complexes were formed based on the theoretical charge ratio (CR) of positive charges from peptide in excess to the negative charges from the pDNA backbone. The optimal charge ratio through experiments was between CR2 and 3 (W/V ratios in Supplementary methods 1). Commercial reagents were used according to manufacturer’s recommendations for transfection of adherent cells. For suspension cells a defined pDNA dose was used for better comparison. For specifications of complex mixing refer to Supplementary methods 1.

As a side note, it should be stated that all hereby presented short term transfection measurements (i.e. luc and EGFP reporters) have been conducted without any means of retaining the transfection-positivity in daughter cells. On the other hand, all the protein production experiments have been conducted by using the QMCF technology that includes cell lines with EBNA/PyLT elements and transfection with compatible plasmid expression vectors.

### 3.4 Firefly luciferase reporter protein expression detection from cell lysate

For reporter luminescence (firefly luciferase expressed from pLuc) assessment in adherent cell culture 10,000 cells per 96 well-plate well were seeded in 100 μl of media one day before transfection. Shortly prior transfection, cell media was replaced with 100 μl of serum free DMEM media. Cells were transfected with 0.2 μg of pLuc per well (refer to 3.3). 4 h post-transfection media was replaced with 100 μl of serum containing media. 24 h post-transfection, media was aspirated, cells washed with 1 x PBS and 30 μl of lysis buffer (0.1% Triton X100 in 1 x PBS). was added For experiments with suspension cells 30,000-35,000 cells per 96 well-plate well in 100 μl of serum free media were seeded 1 h prior transfection. Cells were transfected with 0.1 μg of pLuc per well. 4 h post-transfection 100 μl of fresh serum free media was added. 48 h post-transfection from the final volume of 200 μl from top ν of volume was removed after cell sedimentation and to the rest, 50 μl of lysis buffer was added (0.1% Triton X100 in 1 x PBS). For both, following 20 min incubation at 4°C for cell lysis, the lysate was mixed and 20 μl transferred to a black frame white well 96-well plate for luminescence measurement after addition of substrate in buffer (refer to Supplementary 2). Luminescence signal was detected with GLOMAX 96 microplate luminometer equipped with GLOMAX 1.9.2 software (Promega). RLU values were converted to RLU/mg by normalization to total protein in cell lysate. For protein detection PierceTM BCA Protein Assay Kit was used. Briefly, absorbance of samples was measured at 562 nm after incubation of samples with detection mix.

### 3.5 Secreted alkaline phosphatase expression detection from cell media

For luminescence (SEAP, secreted alkaline phosphatase from pSEAP) detection 600,000-750,000 suspension CHO-K1 or HEK293FT cells per 24-well plate well were seeded 1 h prior experiment in 500 μl of media. Cells were transfected in serum free media with 0.75 μg of pSEAP; 4 h post-transfection 1:1 volume of fresh serum free media was added to each well. 48 h post-transfection 200 μl of cell suspension was collected to a microcentrifuge tube. Tubes were centrifuged (500 x g, 5 min) to pellet the cells. Supernatant was collected and heated for 30 min at 65°C, then cooled to room temperature. 25 μl of sample was transferred to black frame white well 96-well plates for luminescence measurement. To each sample 50 μl of HEPES buffer (10 mM, pH 7.4) and 25 μl of diluted (1 mM) CDP-Star™ substrate was added. Luminescence was measured over 25 min period until the signals equilibrated. Luminescence signal was detected with GLOMAX 96 microplate luminometer equipped with GLOMAX 1.9.2 software (Promega). Results are shown as relative luminescence units (RLU).

### 3.6 Green fluorescent protein (GFP) -expressing cell population detection with flow cytometry

For flow cytometry experiments detecting transfected cell population (green fluorescent protein expression from pGFP) from adherent cells, 50,000 CHO-K1 or HEK293 cells per well were seeded in 500 μl of serum containing media on a 24-well plate 24 h prior transfection. Shortly prior transfection, media was replaced with 500 μl of fresh serum free media. Cells were transfected with 0.5 μg pDNA per well. 4 h post-transfection media on cells was replaced with 500 μl of serum containing media. 24 h post-transfection media was removed, cells washed with 1 x PBS and 0.25% trypsin-EDTA was added to detach cells from the plate. After detachment 1 x PBS supplemented with 1% FBS was added, and cells were analyzed by flow cytometry do detect cell population and fluorescence positive cells. For suspension CHO-K1 and HEK293FT cells 1 h prior experiment 600,000-750,000 cells per 24-well plate well were seeded in 500 μl of serum free media, and transfected with 0.75 μg of pDNA per well. 4 h post-transfection 1:1 volume of fresh serum free media was added to each well. For suspension cells a 200 μl sample was collected 48 h post-transfection for analysis. For flow cytometry sanalysis Attune™ NxT Flow Cytometer equipped with Attune™ NxT Software 3.2.1 was used. For gating, the forward-scatter (FSC) and side-scatter (SSC) of untreated cells were used. Events with high (clumped cells) or low (debris, complexes, media components) size were excluded. Untreated cells were used to set the GFP+ threshold (∼1.5% of untreated were gated as GFP+). For detection 488 laser with 515-545 nm filter was used.

### 3.7 Live/Dead cell viability assay: flow cytometry

For flow cytometry Live/Dead assay (Calcein AM and Propidium iodide-PI detection) in adherent cells 50,000 CHO-K1 or 75,000 HEK293 cells were seeded per 24-well plate well in serum containing media 24 h prior transfection. Cells were transfected with 0.5 μg Firefly luciferase encoding (pLuc) pDNA per well in 500 μl of serum free media; 4 h post-transfection media was replaced with 500 μl of serum containing media. 24 h post-transfection media was removed, cells washed with 1 x PBS and 0.05% trypsin-EDTA was added to detach cells from the plate. Calcein AM (0.4 μM on cells) and PI (1 μl per well) diluted in 1 x PBS were added to the cell suspension 15 min prior analysis. For suspension CHO-K1 and HEK293FT cells 1 h prior experiment 600,000-750,000 cells per 24-well plate well were seeded in 500 μl of serum free media, and transfected with 0.75 μg of pSEAP per well. 4 h post-transfection 1:1 volume of fresh serum free media was added. For suspension cells 200 μl samples were collected from cells 48 h post-transfection. Samples were centrifuged (500 x g, 5 min) and supernatant discarded. The cell pellet was suspended in 250 μl of detection mix consisting of Calcein AM (2 μl of 4 mM stock) and propidium iodide (20 μl) in 14 ml of 1 x PBS. After 15 min incubation at room temperature cells were analyzed. For both, the forward-scatter (FSC) and side-scatter (SSC) of untreated cells were used for gating cell population. Events with high (clumped cells) or low (debris, complexes, media components) size were excluded. The gating for Live/Dead cell population signals from untreated cells, cells incubated with either Calcein AM or PI, and cells pretreated with Triton X100 were used. For Calcein AM 488 laser with 515-545 nm filter was used, and for PI 561 nm laser with 577-593 nm filter was used.

### 3.8 Live/Dead cell viability assay: confocal microscopy

For confocal microscopy 25,000 adherent CHO-K1 cells were seeded on 8-well Nunc™ Lab-Tek™ (Thermo Scientific™, United States) and incubated overnight. Media was replaced with serum free media and cells transfected with 0.25 μg of pLuc plasmid. 24 h post-transfection cells were washed and media replaced with phenol red free DMEM. Confocal images were acquired from live cells with Zeiss LSM710 (Carl Zeiss AG, Germany). For detecting live cells (Calcein AM), 488 nm laser with 593-590 filter was used, for detecting dead cells (PI), 561 nm laser with 593-712 filter was used. The images were taken with 20x magnification. For analyzis Zen software was used.

### 3.9 Trastuzumab mAb expression

For Trastuzumab monoclonal antibody expression 3,000,000 CHO 1E9 cells in Xell CHO TF or 293ALL cells in Balan CD HEK293 media were seeded on a 6 well plate and 2 ml of media shortly prior transfection. For transfection 2 μg of Trastuzumab monoclonal antibody expressing pDNA per well was used. For CHO cells 1 day post-transfection 6% of Xell Basic Feed was added. 3rd day post-transfection 10% of Xell Basic feeed was added and cells transfecred to 30°C incubator. 6th day post-transfection 10% of Xell Basic feed was added and on 8th day post-transfection experiment was terminated. For HEK cells 1 day post-transfection 20% of Xell HEK TF media was added. On 4th day post-transfection 6% of Xell HEK TF was added and on 5th day post-transfection experiment was terminated.

### 3.9 Statistical analysis

Statistical analyses of cell culture experiments were done on the GraphPad Prism software 5.0 (Graphpad Software, CA, USA). All results are shown as a mean with the SEM of at least three separate experiments, if not indicated otherwise. Pearson correlations and 1-way ANOVA was performed with the Statistica application (Dell)

## 4. Conclusions

We were most interested in uncovering connections that could predict the long term and large-scale protein production. Such data would support the future research towards rapid and high throughput lab assays that can be used for the development of new and efficient transfection methods. In the current report we screened a number of various methods that reflect different aspects of transfection efficacies and toxicities in two producer cell lines and we constructed a correlation matrix between the numerical outputs of all the assays for both the cell lines. The correlation matrix is presented in Supplementary Table 3.

When combining all the above data, the most significant conclusions from the above are as follows. First, in CHO, the protein production yields correlated with both suspension expression of SEAP (r=0.88, p < .05) and GFP+ population (r=0.86, p < .05), but did not correlate with luciferase quantitation neither in adherent nor suspension media (Supplementary Table 3,Fig S 14). Although suspension culturing is an important aspect in cell factories, total luciferase reporter levels were not essential in the suspension culture, and the same applies in HEK293 cell line, forcing us to conclude a surprising implication: luciferase reporter quantitation assay, despite its excellent technical aspects, is not a good screening assay for predicting efficacies in protein expression.

The second implication is, as clearly illustrated in the correlation analysis (Supplementary Table 3), the adherent culture methods generally fail to predict the effects in suspension conditions. Neither transfection efficacy assays, nor viability/toxicity assays correlate with protein production efficacies in CHO and HEK293. Hence, when working with the applications of protein expression in mammalian cells, suspension culturing is a must, even in research laboratory settings.

Third, while the current report exemplifies a number of useful predictors for the CHO producer cell line in terms of efficacy assays and transfection methods, the utility and implications for the HEK293 is unfortunately less significant. Although we currently demonstrated that a CPP-based transfection with NF71 significantly outperformed polyplex and lipoplex methods in protein production in HEK293, predictive transfection assays for this cell line should be explored in the future.

Finally, in the current report, we presented several cell penetrating peptide -based transient transfection methods that significantly outperformed widely used method, the polyethylene imine, as well as other polyplex and lipoplex methods. We showed that NF55 and NF51 are efficient for the CHO-based production, and NF71 is an excellent performer in HEK293 cells. These methods have a potential to replace PEI in the industrial settings and offer higher yields for the therapeutic protein production.

## Supplementary data

### Supplementary Methods

**Supplementary methods 1**. Complex mixing specifications per 1 μg of pDNA. Describes mixing ratios for complexes formed between pDNA and transfection reagents.

Supplementary methods 2. **Total Firefly luciferase quantitation and whole cell lysate protein concentration assay**

Describes firefly luciferase measurement assay components, method setup and measurement equipment, and how the results are convereted.

Supplementary methods 3. **MTS-based proliferation assay in adherent cell culture**

Describes the MTS proliferation assay experimental setup.

**Supplementary methods 4**. BrdU (de novo DNA synthesis) proliferation assay Describes the BrdU proliferation assay experimental setup.

### Supplementary Tables

Supplementary table 1. Cell-penetrating peptides used in this work and their sequences. Shows the sequences of the CPPs, references added.

### Supplementary Figures

Supplementary figure 1. **Optimization of complex formation conditions in adherent CHO-K1 and HEK293 cells assessed by total reporter luciferase quantitation from cell lysate**

Shows the results from adherent cell experiments where luciferase is quantified. Different ratios of transfection reagent were tested.

Supplementary figure 2. **The pDNA dose and media change effect based on total reporter luciferase quantitation from cell lysate of transfected adherent CHO-K1 and HEK293**

Shows the results from adherent cell experiments where luciferase is quantified. Different pDNA doses were tested, and if the media change had a considerable effect on the RLU/mg.

Supplementary figure 3. **Optimization of suspension cell culture transfection volume and time assessed by total reporter luciferase quantitation**

Shows the results from suspension cell experiments where luciferase is quantified. Analysis time-point 24 h and 48 h were compared, and the transfection media volume was tested.

Supplementary figure 4. **Optimization of suspension CHO-K1 and HEK293FT cell transfection conditions assessed by total reporter luciferase quantitation**

Shows the results from suspension cell experiments where luciferase is quantified. Different ratios of transfection reagent were tested.

Supplementary figure 5. **Fluorescence intensities of transfected adherent cell population expressing green fluorescent protein**

Shows the results from flow cytometry with suspension cells. The fluorescence is divided by intensities of the signal.

Supplementary figure 6. **Transfected suspension cell population expressing green fluorescent reporter protein**

Results from flow cytometry with suspension cells. Showing the fluorescence signal over a 5-days post-transfection. The population with fluorescence signal measured 48 h post-transfection is higher than at 24 h.

Supplementary figure 7. **Flow cytometry profiles of transfected suspension CHO-K1 cells expressing green fluorescent protein**

Results from flow cytometry with suspension cells. The profiles of fluorescence intensity in CHO-K1 cells 48 h post-transfection.

Supplementary figure 8 **Flow cytometry profiles of transfected suspension HEK293FT cells expressing green fluorescent protein categorized to low, medium and high expressing cell populations**.

Results from flow cytometry with suspension cells. The profiles of fluorescence intensity in HEK293FT cells 48 h post-transfection.

**Supplementary figure 9**. Proliferation of cells post-transfection assessed by metabolic activity (MTS) and de novo synthesis assessment (brdU) in adherent CHO-K1 and HEK293 cells.

Results from proliferation assays with adherent cells.

**Supplementary figure 10**. Live/Dead assay control groups in adherent CHO K1 and HEK293 Results of control groups from flow cytometry with suspension cells.

**Supplementary figure 11**. Live/Dead assay of suspension CHO-K1 and HEK293FT cells analyzed 24 h and 48 h post-transfection Results from flow cytometry of Live/dead assay with suspension cells 24 h post-transfection.

**Supplementary figure 12**. Live/Dead assay of suspension HEK293FT cells analyzed 24 h and 48 h post-transfection

Results from flow cytometry of Live/dead assay with suspension cells 24 h, and 48h post-transfection.

**Supplementary figure 13**. Gel images for Figure 3

**Supplementary figure 14**. Correlation analysis graphs

**Supplementary methods 1. Transfection complex mixing specifications per 1 μg of pDNA**.

For NF51 the ratio per 1 μg of pDNA was defined as 1.5 μl (CR2) or 2.3 μl (CR3) of 1 mM peptide. For NF71 the ratio per 1 μg of pDNA was defined as 1.3 μl (CR2) or 2 μl (CR3) of 1 mM peptide. For PF14 the ratio per 1 μg of pDNA was defined as 1.2 μl (CR2) or 1.8 μl (CR3) of 1 mM peptide. For NF55 the ratio per 1 μg of pDNA was defined as 2 μl (CR2) or 3 μl (CR3) of 1 mM peptide. The complex was formed in 1/10^th^ of final transfection volume. The volume was adjusted by adding ultrapure water to pDNA prior addition of peptide to the complex mix. Complexes were incubated 20-40 minat room temperature before addition to cells.

For PEI the N/P ratio is used to calculate complex formation ratios. N/P ratio reflects ratio of polymer amine (N or nitrogen) groups to nucleic acid phosphate (P) groups. The reagent was diluted in milliQ water to concentration 1 mg/ml and added to pDNA diluted in milliQ water. The mix was vortexed for 5 s and incubated at room temperature for 20 min prior addition to cells. For N/P 20 2.6 μl, N/P 30 3.9 μl and N/P 60 7.8 μl of 1 mg/ml PEI was added per 1 μg of pDNA in ultrapure water. For Reagent007 the ratio per 1 μg of pDNA was defined as 4.75 μl of working solution mixed in ultrapure water. Complexes were incubated 5 min at room temperature prior addition to cells. For Lipofectamine3000 the ratio per 1 μg of pDNA was defined as 2 μl of P3000 and 1.5 (low) – 3 (high) μl of LF3000 reagent. For complex formation pDNA and P3000 were mixed in serum free media (mix A). LF3000 was diluted in serum free media (mix B). Both components (A+B) were then mixed, suspended and incubated at room temperature 10 min prior addition to cells. For Turbofect the ratio per 1 μg of pDNA was defined as 1-2 μl (2 μl used in this work) of reagent mixed in serum free media. Plasmid was mixed with serum free media, and then reagent was added. Complexes were incubated at room temperature for 20 min prior addition to cells. For Xfect the ratio per 1 μg of pDNA was defined as 0.3 μl of reagent, mixed in reaction buffer. Plasmid was mixed with buffer, vortex for 5 s, the reagent was added, vortexed for 10 s and complexes were incubated at room temperature for 10 min prior addition to cells. For jetOPTIMUS the ratio per 1 μg of pDNA was defined as 1 μl of reagent, mixed in reaction buffer. Plasmid was mixed with buffer, vortexed, reagent added, vortexed and incubated at room temperature for 10 min prior addition to cells. For TransIT X2 the ratio per 1 μg of pDNA was defined as 2-6 μl of reagent (3 μl used in this work). Plasmid was mixed with serum free media, and then reagent was added. The complex solution was mixed by pipetting and incubated for 20 at room temperature prior addition to cells.

**Supplementary methods 2. Total Firefly luciferase quantitation and whole cell lysate protein concentration assay**

Firefly luciferase was detected from the cell lysates of cells transfected with 0.1-0.2 μg of Firefly luciferase encoding pLuc. For adherent CHO-K1 and HEK293 cells 10,000 cells per well were seeded in 100 μl of serum containing media on a 96-well plate 24 h prior transfection. Briefly prior transfection the media was replaced with 100 μl of serum free DMEM media. 4 h post-transfection media was replaced with serum containing media (if not stated otherwise), and further incubated. 24 h post-transfection the media was aspirated, cells washed with 1 x PBS and lysis buffer added to wells. Lysis buffer consisted of 0.1% Triton X100 in 1 x PBS buffer. After 20 min incubation, the lysate was mixed and used for further analysis.

For suspension CHO-K1 and HEK293FT cells 30,000-35,000 cells per well were seeded in 100 μl of serum free media on a 96-well plate ∼1 h prior transfection. For CHO-K1 cells Xell CHO TF media was used, and for HEK293FT cells Gibco Freestyle293 media was used. Both medias were supplemented with GlutaMAX. 4 h post-transfection 100 μl of fresh serum free media was added to each well. 24-48 h post-transfection cells were let to sediment and ½ of media was removed from the top. To the rest 50 μl of lysis buffer was added and cells incubated for 20 min for lysis. Lysate was mixed and used for further analysis.

Luminescence from expressed firefly luciferase was detected by mixing 20 μl of cell lysate with 100 μl of firefly luciferase detection solution. The detection solution was prepared shortly prior addition to cells, by mixing DTT (25 nM final concentration), D-Luciferin (1 mM), ATP (1 mM), Coenzyme A (25 μM), EDTA (1 mM), Tricine (20 mM), MgCO_3_ (1 mM) in ultrapure water. After addition of solution, the signal (relative luminescence units, RLU) was detected with GLOMAX 96 microplate luminometer equipped with GLOMAX 1.9.2 software (Promega).

To determine total protein cincentrations from the cell lysate PierceTM BCA Protein Assay Kit was used. Cell lysate was transferred to a transparent 96-well plate and to each sample 100 μl of solution consisting of component A and B mixed in 50:1 ratio was added. Following 30 min incubation absorbance was measured at 562 nm wavelength. Absorption values were converted to protein concentrations using standard curve generated by measuring samples with known protein concentrations. Absorption was measured with Tecan Sunrise microplate absorbance reader (Tecan Group Ltd, Switzerland). RLU values were converted to RLU/mg by normalization to protein in the cell lysate.

**Supplementary methods 3. MTS-based proliferation assay in adherent cell culture**

10,000 CHO-K1 and HEK293 cells per well were seeded 24 h prior transfection on a 96-well plate in 100 μl of serum containing phenol free DMEM media supplemented with 1mM sodium pyruvate, 0.1Mm non-essential amino acids, 100 U/mL penicillin, and 100mg/mL streptomycin. Shortly prior transfection media on cells was replaced to 100 μl of serum free media and complexes containing 0.2 μg of pLuc pDNA were added to each well. CellTiter 96® AQ_ueous_ One Solution Cell Proliferation Assay (MTS) was used as indicated by the manufacturer’s instructions. 20 h post-transfection to each well 20 μl CellTiter 96® AQueous One Solution Reagent was added and cells further incubated for 4 h in a humidified, 5% CO_2_ atmosphere at 37°C. Following incubation, absorbance was measured at 490 nm using a Tecan Sunrise microplate absorbance reader (Tecan Group Ltd, Switzerland) to detect soluble formazan produced by cellular reduction of MTS.

**Supplementary methods 4. BrdU (de novo DNA synthesis) proliferation assay**

10,000 CHO-K1 and HEK293 cells were seeded 24 h prior transfection on a 96-well plate in 100 μl of serum containing phenol free DMEM media supplemented with 1mM sodium pyruvate, 0.1mM non-essential amino acids, 100 U/mL penicillin, and 100mg/mL streptomycin. Shortly prior transfection media was replaced with 100 μl of serum free media and complexes containing 0.2 μg of pLuc were added to each well. 4 h post-transfection media was replaced with serum containing media and BrdU reagent was added.

Abcam brdU Cell Proliferation ELISA kit (colorimetric) kit was used as indicated in the manufacturer’s instructions. Bromodeoxyuridine / 5-bromo-2’-deoxyuridine is an analog of the nucleoside thymidine used in the BrdU assay to identify proliferating cells. 24 h post-transfection the media was removed, 200 μl of Fixing solution was added and cells incubated at room temperature for 30 min. Fixing solution was aspirated and cells washed three times with 200 μl of wash buffer. To each well 100 μl of anti-brdU monoclonal antibody solution was added and incubated with cells at room temperature for 1 h. Antibody solution was removed, and the cells were washed three times with 200 μl of wash buffer. To each well 100 μl of 1 x peroxidase goat anti-mouse IgG conjugate solution was added and incubated at room temperature for 30 min. The solution was then aspirated, and cells washed three times with 200 μl of wash buffer. Then the plate was flooded with ultrapure water and then tapped dry. To each well 100 μl of TMB peroxidase substrate was added and incubated in dark, at room temperature for 30 min. Reaction was then stopped with 100 μl of stop solution. Absorbance was measured at 450/550 nm wavelengths with Tecan Sunrise microplate absorbance reader (Tecan Group Ltd, Switzerland). Cells actively proliferating turn blue which is proportional to the brdU incorporation in the proliferating cells.

**Supplementary Table 1.** Cell-penetrating peptides used in this work and their sequences.

| Name | Sequence | Reference |
| --- | --- | --- |
| NickFect51 | Stearoyl-AGYLLGO<sup>a</sup>INLKALAALAKKIL<sup>b</sup> | Arukuusk, et al, 2013, [8] |
| NickFect71 | Stearoyl-HHYHHGO<sup>a</sup>ILLKALKALAKAIL<sup>b</sup> | Porosk, et al, 2019, [11] |
| PepFect14 | Stearoyl-AGYLLGKLLOOLAAAALOOLL<sup>b</sup> | Veiman, et al 2013, [12] |
| NickFect55 | Stearoyl-AGYLLGO<sup>a</sup>INLKALAALAKAIL<sup>b</sup> | Freimann, et al 2016, [9] |

<sup>a</sup> – The synthesis is continued from the sidechain amino group of the amino acid ornithine, instead of alpha-amino group as usually.
<sup>b</sup> – The peptides are C-terminally amidated.

**Fig S 1.**
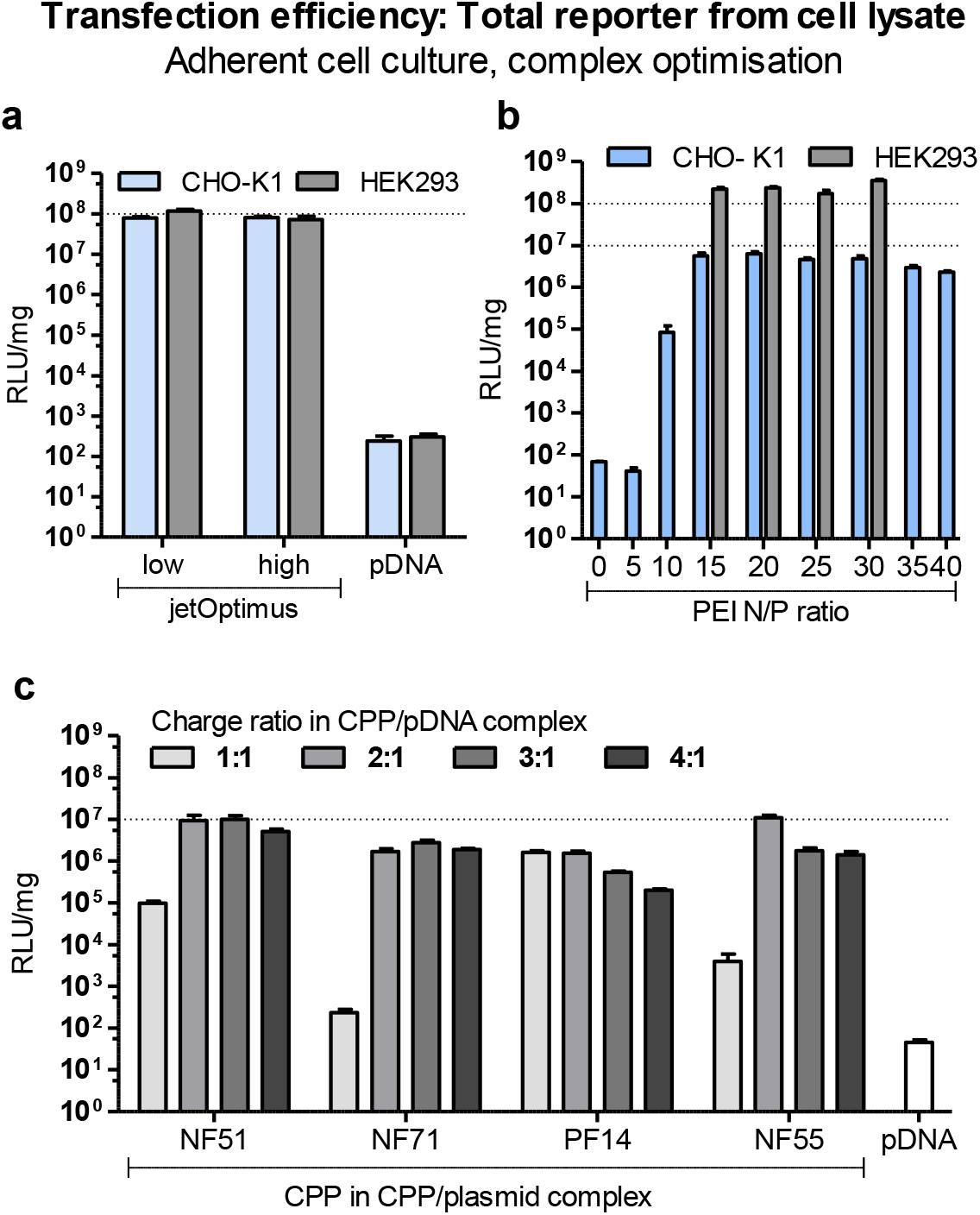
Optimization of complex formation conditions in adherent CHO-K1 and HEK293 cells assessed by total reporter luciferase quantitation from cell lysate. Adherent CHO-K1 or HEK293 cells were transfected with 0.2 μg of Firefly luciferase encoding pLuc per well on a 96-well plate. Transfection was done in serum free DMEM media. Luminescence was measured from the cell lysate 24 h post-transfection and relative luminescence units (RLU) were normalized to protein measured from the lysate. **a**) For jetOPTIMUS the amount of reagent was given as a range of reagent per 1 μg of pDNA. The lowest amount (low) and highest amount (high) of reagent per pDNA dose were tested. **b**) For PEI the optimization range was not indicated within the user protocol, therefore we tested a broader range based on theoretical PEI/pDNA N/P ratios. The N/P 20 was chosen for following experiments in adherent cells. **c**) CPP/pDNA charge ratio screen in adherent cells in serum free media. The charge ratio reflects the ratio between positive charges in the peptide and negative charges in the DNA phosphate backbone. Complexes formed with CPP in excess. On average, depending on the specific experiment, the optimal range was between CR2 and CR3

**Fig S 2.**
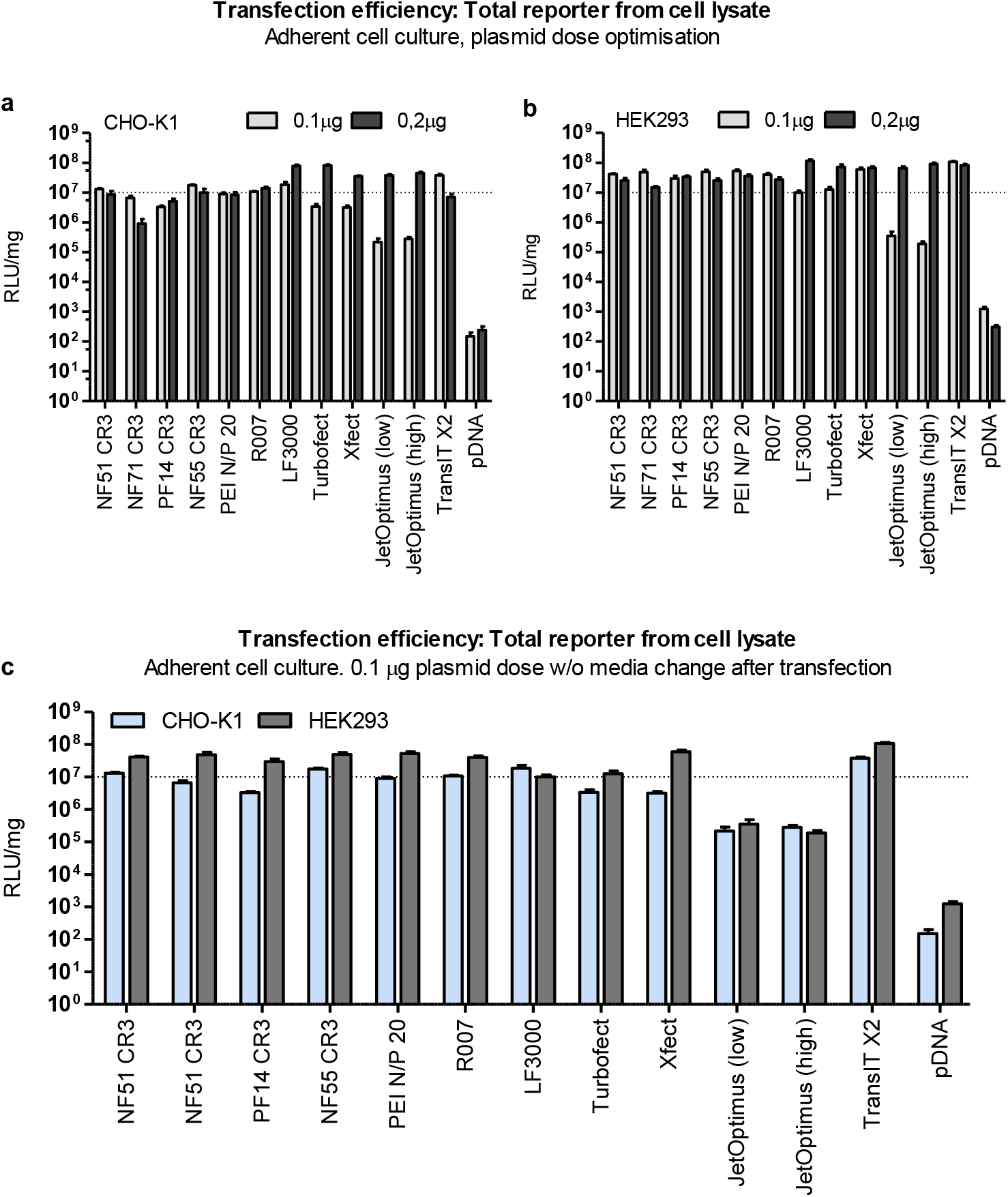
The pDNA dose and media change effect based on total reporter luciferase quantitation from cell lysate of transfected adherent CHO-K1 and HEK293 cells. Adherent CHO-K1 and HEK293 cells were transfected with Firefly luciferase encoding plasmid pLuc. Transfection was done in serum free media. Plasmid dose comparison in a) CHO K1 or b) HEK293 cells. Cells were transfected with either 0.1 μg or 0.2 μg pDNA dose per 96-well plate well in 100 μl of serum free media. 4 h post-transfection media was replaced with serum containing media. 24 h post-transfection cells were lysed and luminescence measured from the cell lysates. Same lysate was used to determine total protein concentration and RLU was normalized to protein in lysate. c) For CPP/pDNA complexes we often use transfection without media change 4 h post-transfection. We compared the transfection efficacy between transfection reagents and CPPs in this setting. The pDNA dose per well was 0.1 μg and transfection was done in serum free media. Media was not changed after addition of complexes, and cells were incubated with complexes in serum free media for 24 h. Following wash and cell lysis, the luminescence and protein concentration were determined from the lysate. Results are expressed as relative luminescence units per mg of total protein (RLU/mg).

**Fig S 3..**
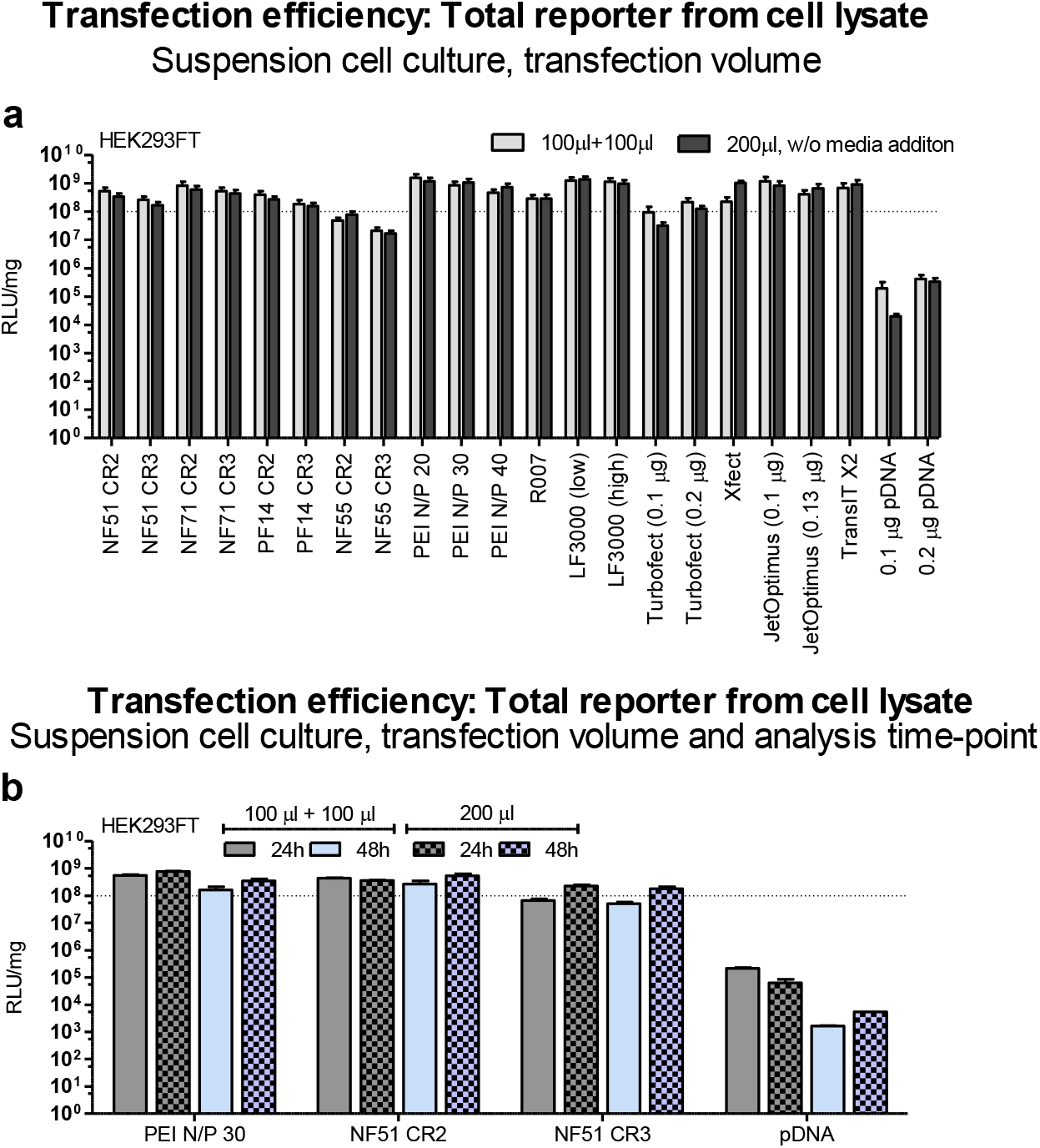
Optimization of suspension cell culture transfection volume and time assessed by total reporter luciferase quantitation. Suspension HEK293FT cells were transfected with 0.1 μg of Firefly luciferase encoding pLuc. To determine whether or not the addition of media would be beneficial to the transfection two setting were compared. With media addition 4 h post-transfection (100 μl + 100 μl), and transfection in the same final volume, but without addition of fresh media post-transfection (200 μl). a) 24 h post-transfection cells were let to sediment and 100 μl of media was removed from the top. To each well 50 μl of lysis buffer (0.1% Triton X100 in 1 x PBS) was added to the remaining 100 μl of media with cells. After lysis 20 μl sample was used to determine RLU. Generally, there were no major differences between transfection efficacies between these two groups. b) Considering the time-point after transfection we screened some groups with the previous settings (100 μl + 100 μl vs 200 μl) at 24 h and 48 h post-transfection. Although the luminescence values for transfected groups did not significantly differ between 24h and 48h measurements, interestingly, the background signal in pDNA treated and untreated groups was lower 48h post-transfection. Compelled by more favorable signal-to-noise ratio we chose 48h post-transfection measurements for suspension cells. For CPP/pDNA complexes CR2 had a slightly higher reporter levels when compared to CR3, therefore we continued with CR2 in suspension cell cultures.

**Fig S 4.**
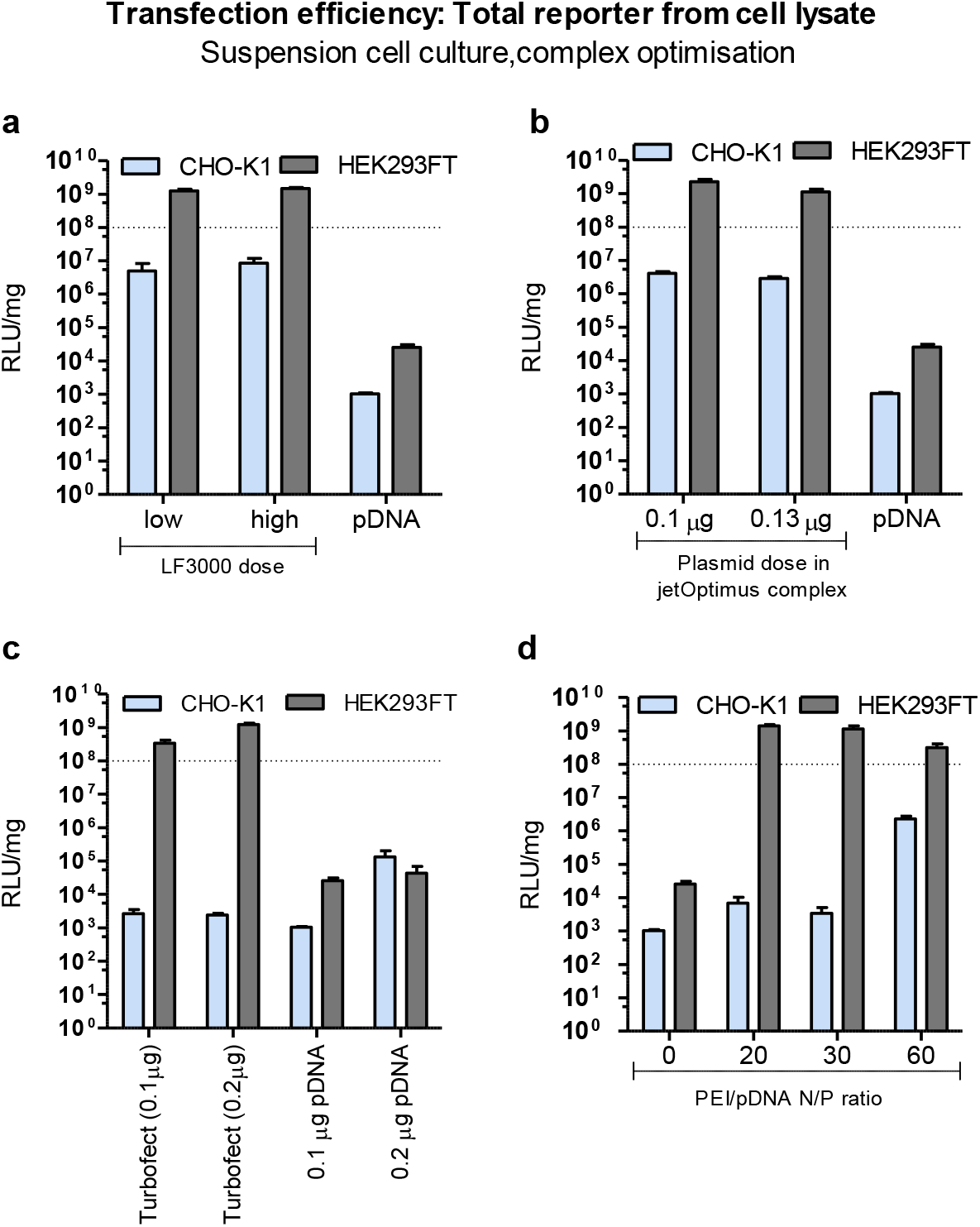
Optimization of suspension CHO-K1 and HEK293FT cell transfection conditions assessed by total reporter luciferase quantitation. Suspension CHO-K1 and HEK293FT cells, transfected with 0.1 μg (if not stated differently) Firefly luciferase encoding pLuc. 4 h post-transfection equal volume of media was added to each well. 48 h post-transfection 100 μl of media was removed, lysis buffer added and from lysate luminescence (RLU) and total protein concentration(mg) were determined. We tested a) LF3000 reagent dose, b) pDNA dose for jetOPTIMUS, c) PEI N/P ratio from 0 to 60, and d) pDNA dose for Turbofect in CHO-K1 and HEK293FT cells. Using higher reagent dose in LF3000/pDNA complexes did not have a significant effect, therefore for further experiments low dose was used in suspension cells. In case of jetOPTIMUS the indicated lowest pDNA dose was 0.13 ug, but also the 0.1 ug pDNA dose did not result in significantly lower signal. In PEI complexes, interestingly for CHO-K1 transfection, the required N/P ratio compared to HEK293FT cells was considerably higher, whereas in HEK293FT cells already the complexes formed at N/P 20 resulted in high reporter expression. In case of Turbofect we also included group with 0.2 μg pDNA dose, as this was indicated minimal amount of pDNA per 96 wp well. The reporter protein levels increased almost 3 fold compared to corresponding pDNA treated group, but even when taking this into account, Turbofect would not be the top performer in the comparison to other transfection reagents tested at 0.1 ug pDNA dose..

**Fig S 5.**
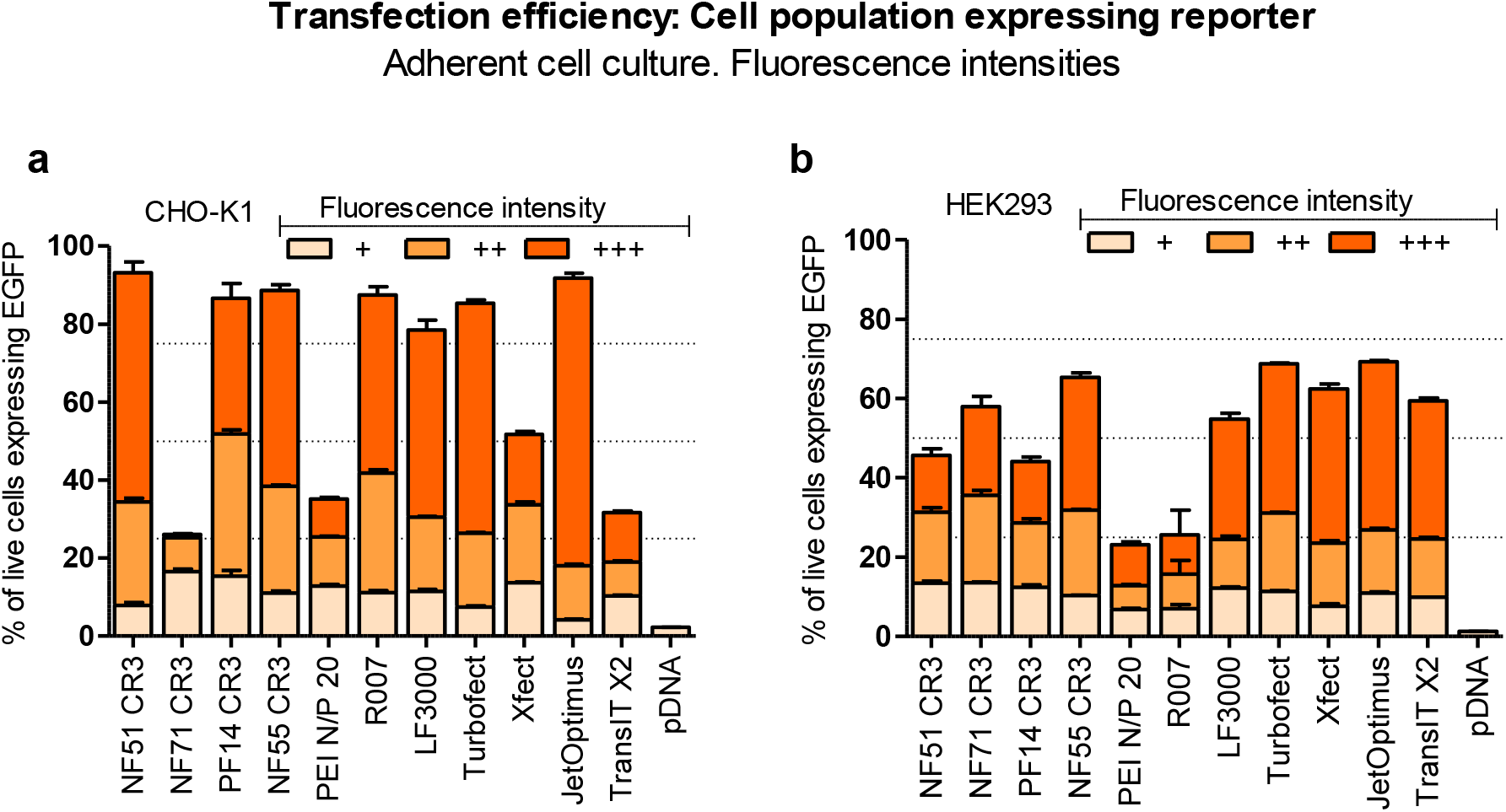
Fluorescence intensities of transfected adherent cell population expressing green fluorescent protein. 24 h prior transfection 50,000 cells per well were seeded on a 24-well plate well in 500 μl of media. Shortly prior transfection media was replaced with serum free media and adherent a) CHO-K1 and b) HEK293 cells were transfected with 0.5 μg of green fluorescent protein expressing pGFP. 4 h post-transfection media was replaced with 500 μl of serum containing media. 24 h post-transfection cells were washed, detached from the plate and cells with fluorescent signal were determined using flow cytometry. For gating untreated cells side-scatter and forward scatter plots were used, and from the cell population the threshold was set to ∼1% fluorescent cells in untreated cell samples.

**Fig S6.**
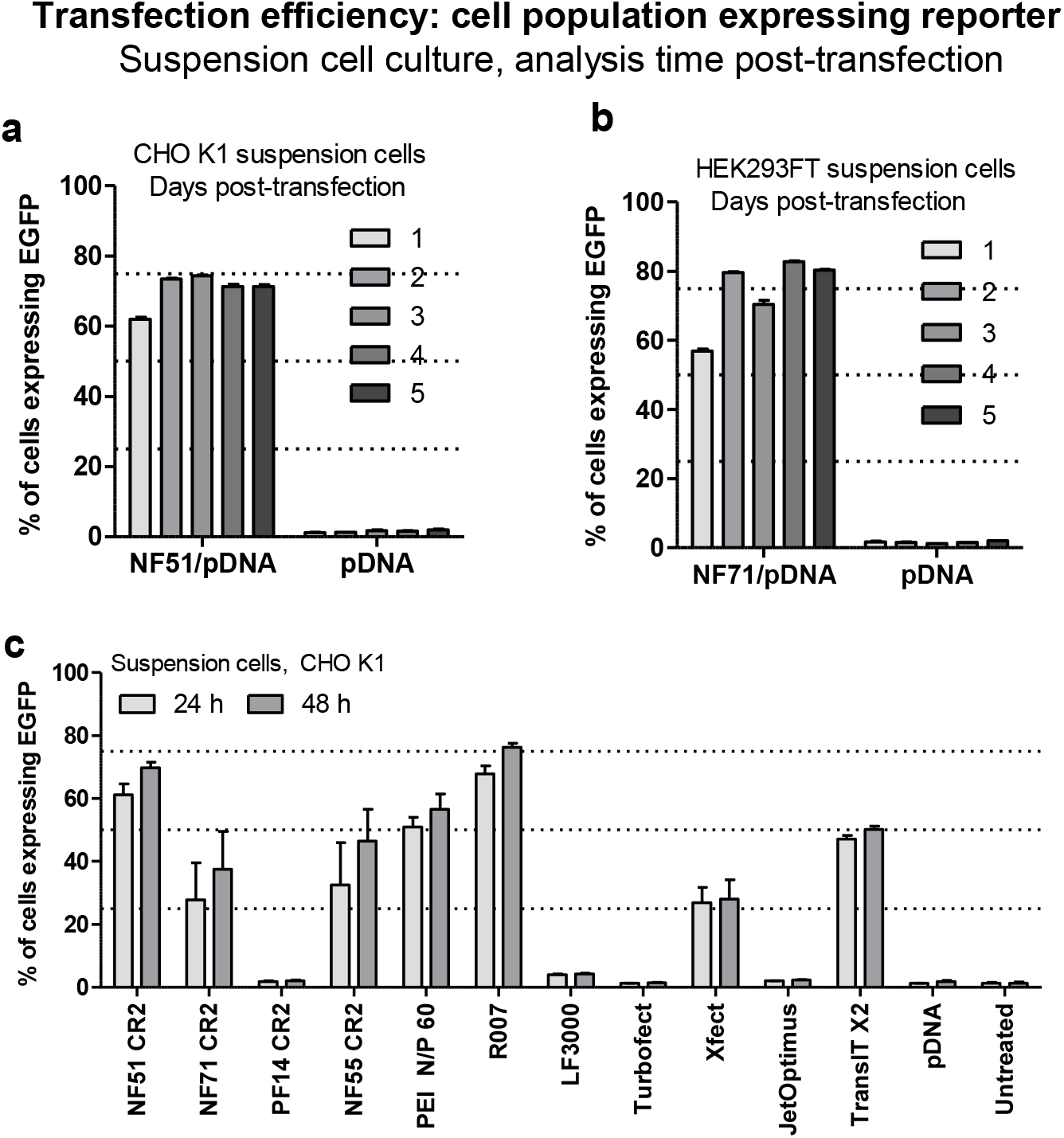
Transfected suspension cell population expressing green fluorescent reporter protein. 1 h prior transfection 600,000-750,000 suspension cells per well were seeded on a 24-well plate in 500 μl of serum free media. For CHO cells Xell CHO TF media was used, and for HEK293FT cells Gibco Freestyle293 media was used. Both media were supplemented with GlutaMAX. The a) CHO-K1 and b) HEK293FT cells were transfected with CPP/pDNA complexes. 0.75 μg of green fluorescent protein encoding plasmid pGFP was used per well. As a control wells with cells treated with pDNA at the same dose were used. Reporter protein (GFP) expressing cell population was detected by flow cytometry 1 to 5 days post-transfection. c) The percentage of green fluorescent protein expressing cell population detected from suspension CHO-K1 cells 24 h and 48 h post-transfection of 0.75 μg of pGFP.

**Fig S 7.**
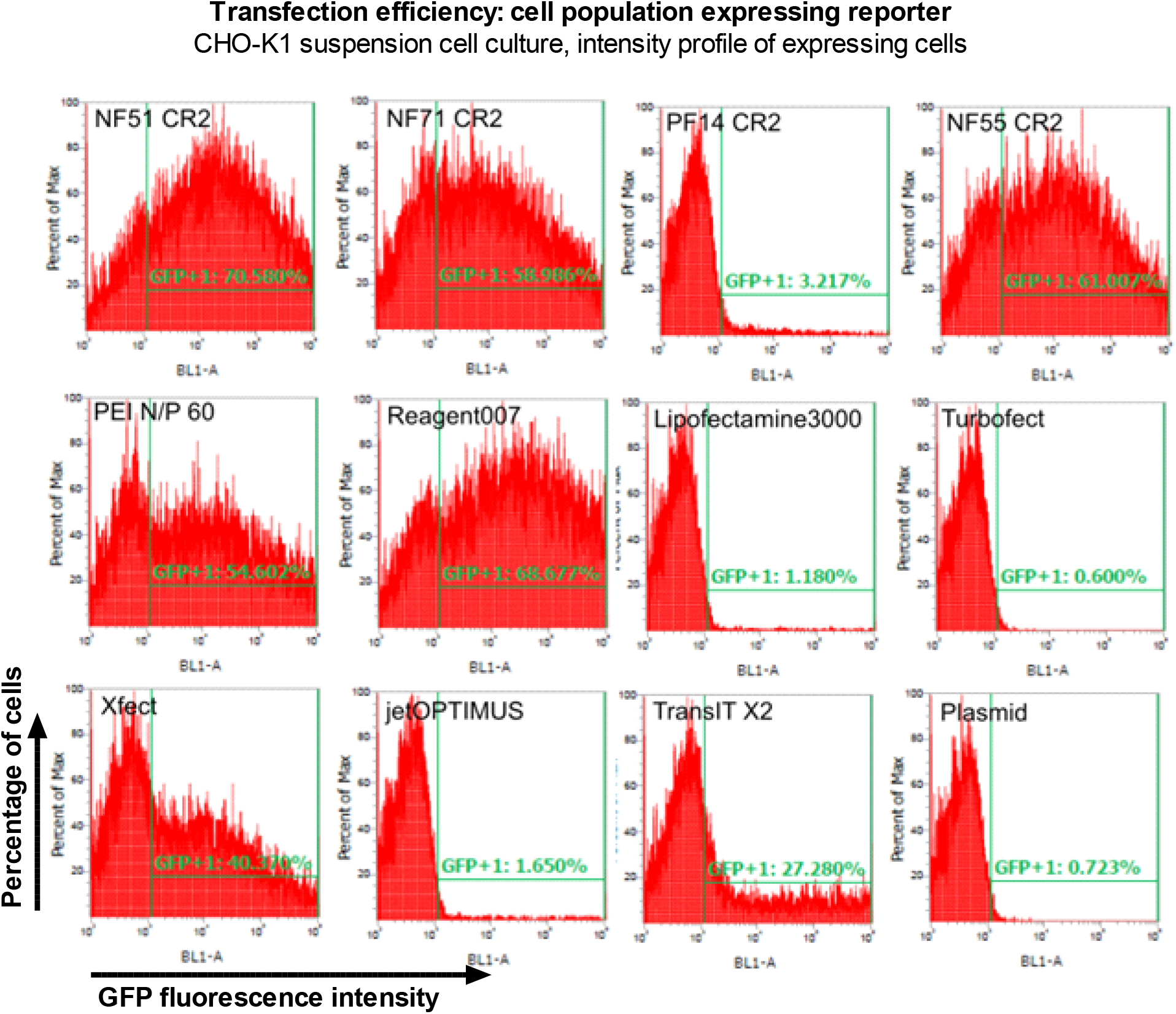
Flow cytometry profiles of transfected suspension CHO-K1 cells expressing green fluorescent protein. 600,000-750,000 suspension CHO-K1 cells per well were seeded on a 24-well plate in 500 μl of serum free Xell CHO TF media 1 h prior transfection. Cells were transfected with 0.75 μg of green fluorescent protein encoding pGFP per well. 4 h post-transfection 500 μl of fresh media was added to cells. 48 h post-transfection cells were analyzed by flow cytometry. The signal intensities from cells and percentages from cell population are shown on each graph. Marked on each is the transfection reagent used for transfection. GFP+ is showing gated events with signal intensity over threshold of ∼1% of untreated cells.

**Fig S 8.**
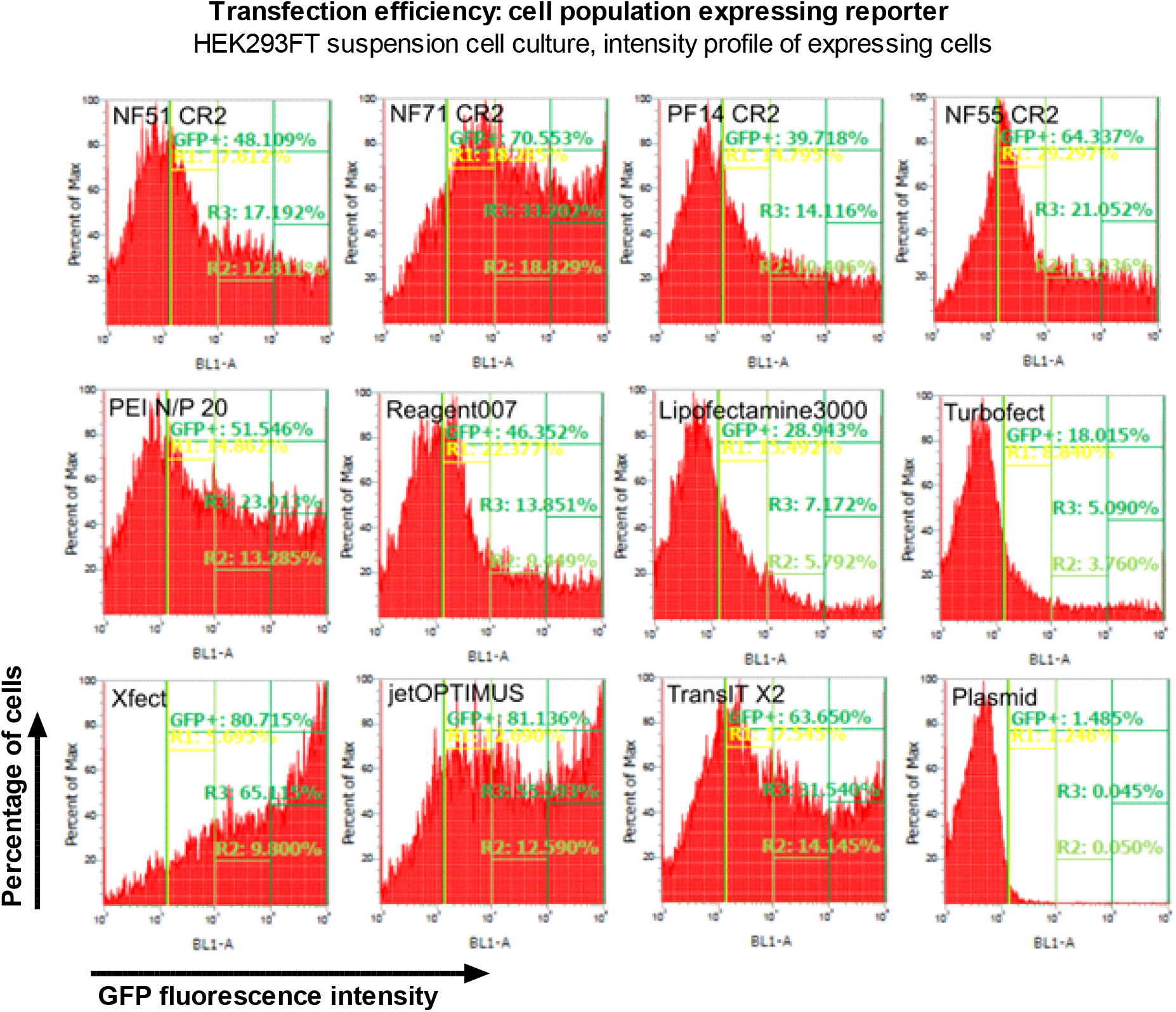
Flow cytometry profiles of transfected suspension HEK293FT cells expressing green fluorescent protein categorized to low, medium and high expressing cell populations. 600,000-750,000 suspension HEK293FT cells per well were seeded on a 24-well plate in 500 μl of serum free Gibco Freestyle293 media 1 h prior transfection. Cells were transfected with 0.75 μg of green fluorescent protein encoding pGFP per well. 4 h post-transfection 500 μl of fresh media was added to cells. 48 h post-transfection cells were analyzed by flow cytometry. The signal intensities from cells and percentages from cell population are shown on each graph. GFP+ is showing gated events with GFP signal.

**Fig S 9.**
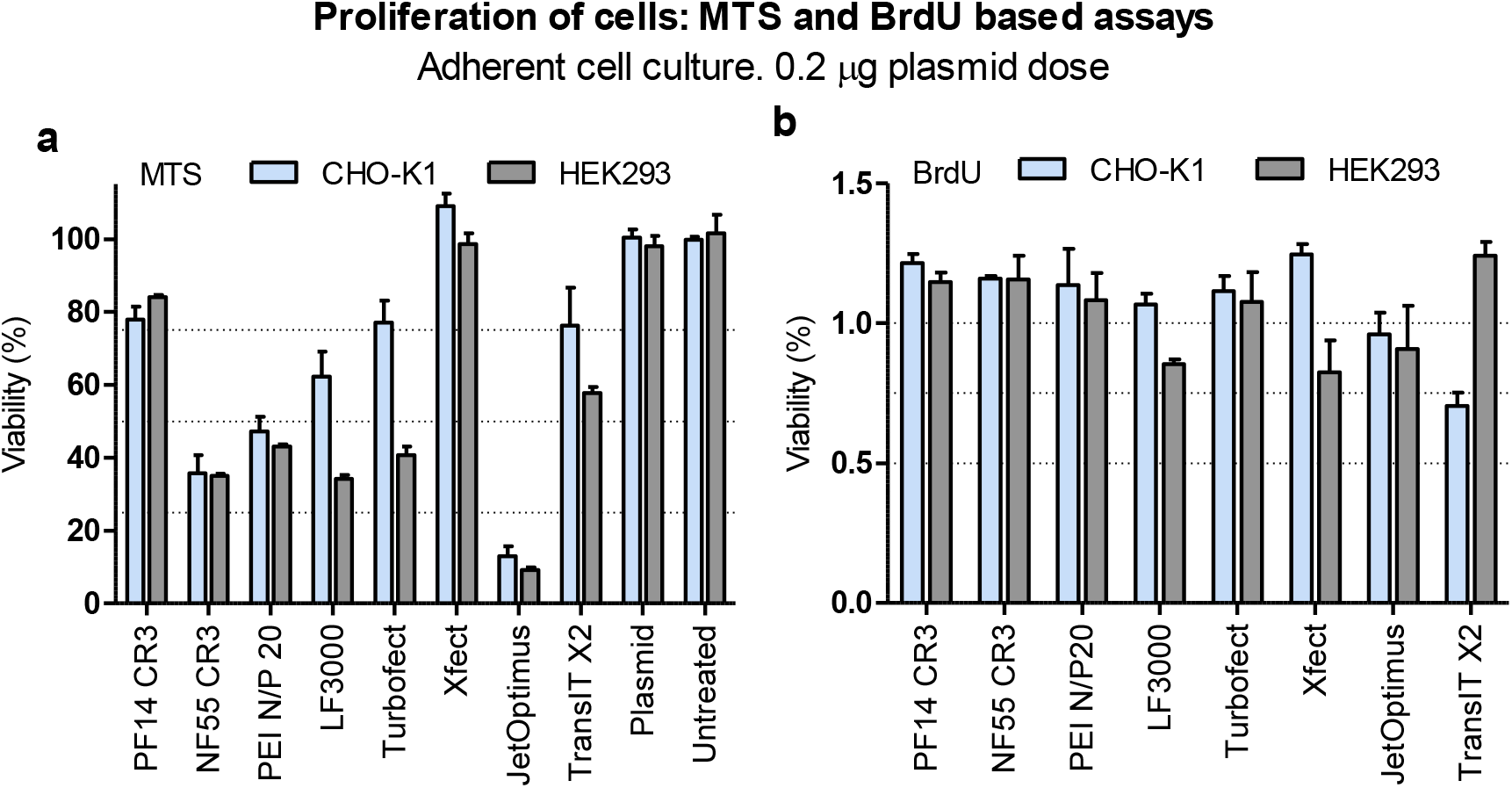
Proliferation of cells post-transfection assessed by metabolic activity (MTS) and de novo synthesis assessment (brdU) in adherent CHO-K1 and HEK293 cells. a) Cells transfected in 100 μl of serum free media with 0.2 μg of Firefly luciferase encoding pLuc per 96-well plate well. 20 h post-transfection 20 μl of CellTiter 96® AQueous One Solution Reagent was added to each well and cells further incubated at humidified incubator for 4 h, following detection of absorbance. Results are shown absorbance values normalized to untreated cells and converted to percentages, where absorbance from untreated cells (100%) and background (0%) was taken into account. (Details in Supplementary methods 3) b) BrdU assay, which reflects the proliferation of cells by incorporation into newly synthesized DNA. Abcam brdU Cell Proliferation ELISA colorimetric kit was used to determine cell proliferation 24 h post-transfection. Results are expressed as fold to untreated (1.0). (Details in Supplementary methods 4). The comparison between MTS and BrdU with the other assays did not yield useful and significant correlations and are not shown in the summary correlation matrix

**Fig S 10.**
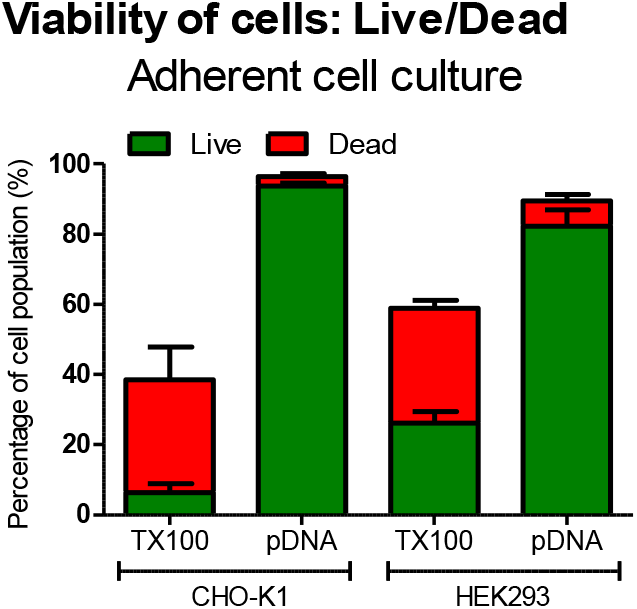
Live/Dead assay control groups in adherent CHO K1 and HEK293. 50,000-a) CHO-K1 and 75,000 b) HEK293 cells seeded 24 h before transfection on a 24-well plate in serum containing DMEM media. Cells were incubated in serum free media with or without 0.5 μg of pDNA pLuc per well. 4 h post-addition of pDNA, the cell media was replaced with serum containing media. 24 h post-transfection cells were washed, detached from the plate and Calcein AM (Live) and PI (Dead) were added. For TX100 group with detection mix 0.1% Triton X100 solution was added to the cells. Flow cytometry was used to detect Live and Dead cells from cell population. Results are expressed as percentage of events from all events with FSC and SSC similar to cellscells.

**Fig S 11.**
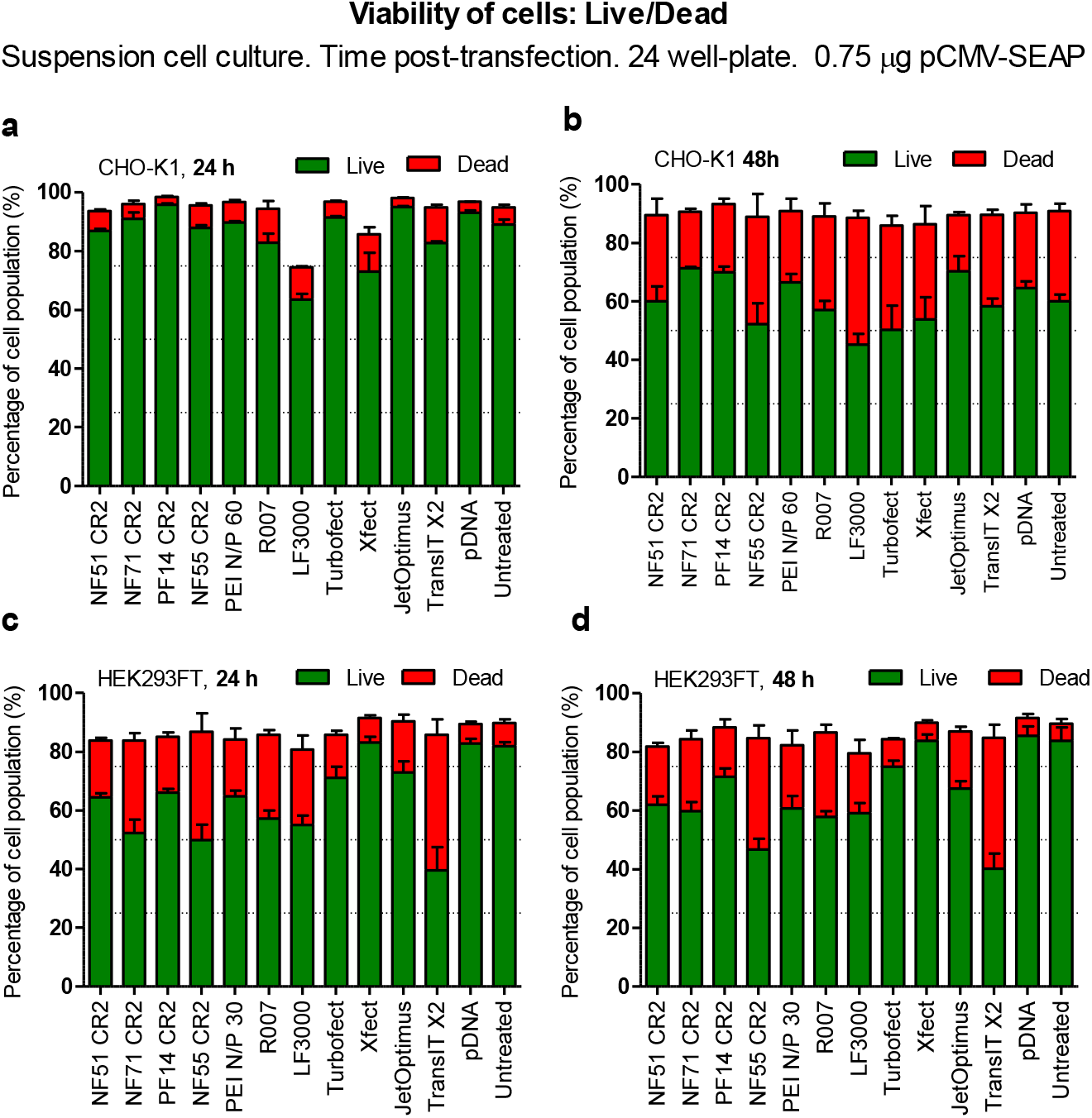
Live/Dead assay on suspension CHO-K1 and HEK293FT cells analyzed 24 h and 48 h post-transfection. 600,000-750,000 suspension a) and b) CHO K1 or c) and d) HEK293FT cells per 24-well plate well were seeded ∼1 h prior transfection in 500 μl of serum free media. Cells were transfected with 0.75 μg of pSEAP per well. 4 h post-transfection 500 μl of fresh media was added. a) and c) 24 h, or b)and d) 48 h post-transfection cells were collected, centrifuged and solution containing Calcein AM and PI was added to detect Live (Calcein AM) and Dead (PI) cells from cell population with flow cytometry

**Fig S 12..**
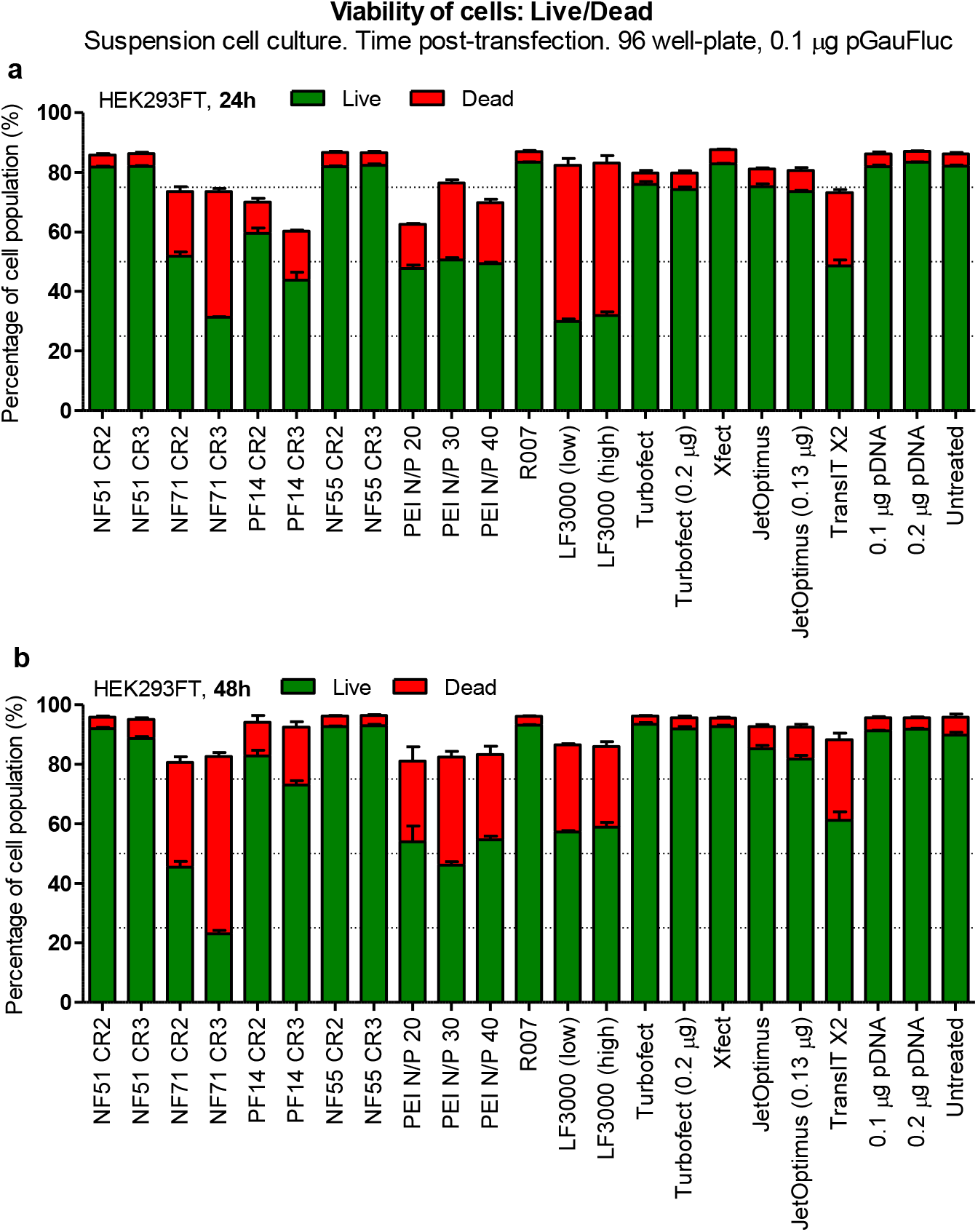
Live/Dead assay on suspension HEK293FT cells analyzed 24 h and 48 h post-transfection. For 96 well plate format 40,000 HEK293FT suspension cells per well were seeded in 100 μl of media per well 1 h prior transfection. Cells were transfected with 0.1 μg of Firelfly luciferase encoding pLuc. 4 h post-transfection 100 μl of fresh media was added. 24 h and 48 h post-transfection from 200 μl cell suspension 100 μl was collected and Calcein AM and PI was added to detect Live (Calcein AM) and Dead (PI) cells from cell population by flow cytometry.

**Fig S 13..**
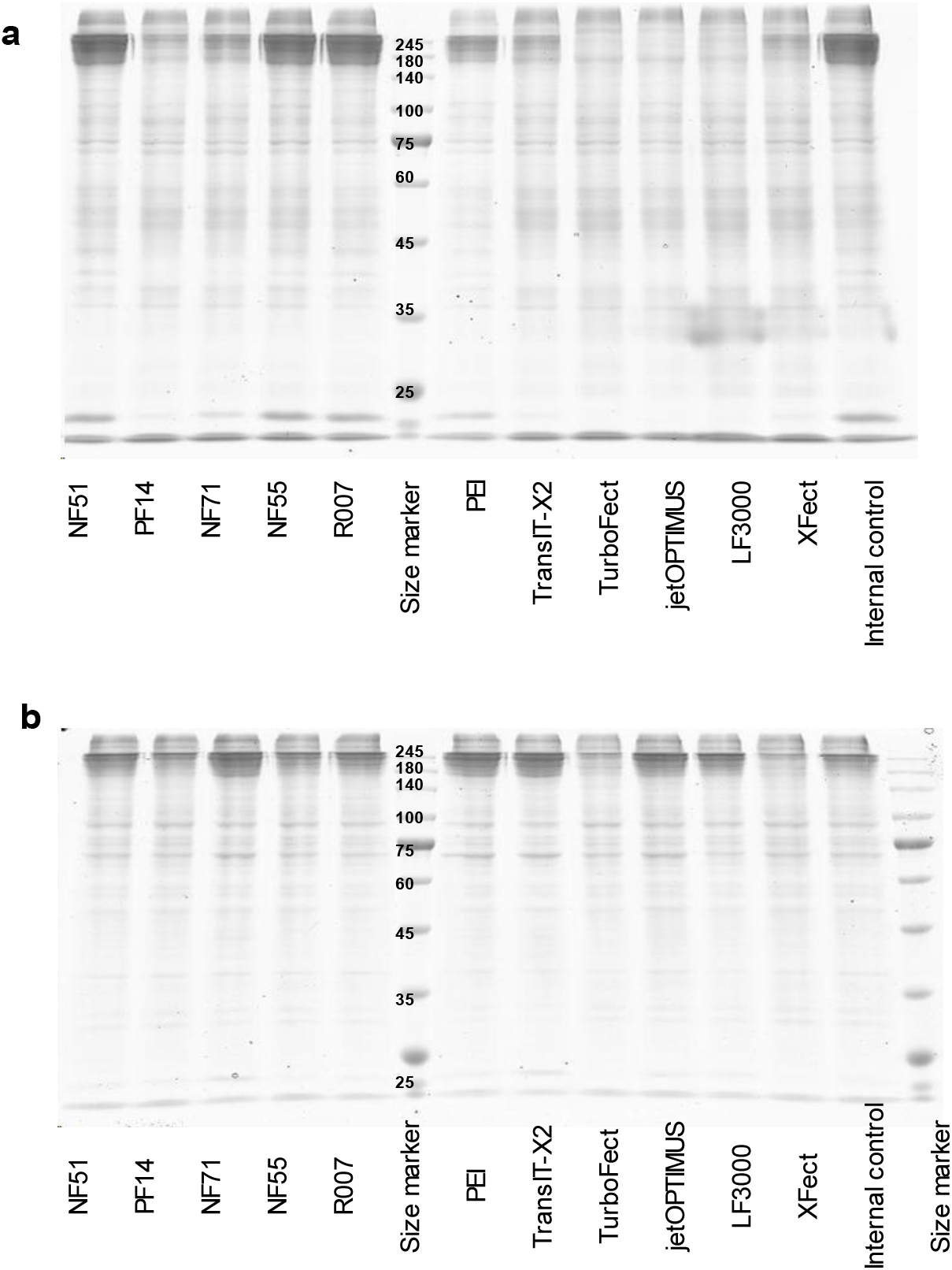
Full SDS gel images used in Figure 3. CHO and HEK cells used to express Trastuzumab mAB in suspension cell culture. 2 μg of pLic2.1 pDNA per 6-well plate well was used. For CHO 1E9 (a) cells transfection in Xell CHO TF serum free media was used and analysis was performed 8 days post-transfection. For HEK293ALL(b) BalanCD HEK293 serum free media was used and analysis was performed 5 days post-transfection. For CPP/pDNA complexes CR2.5 was used

**Fig S 14.**
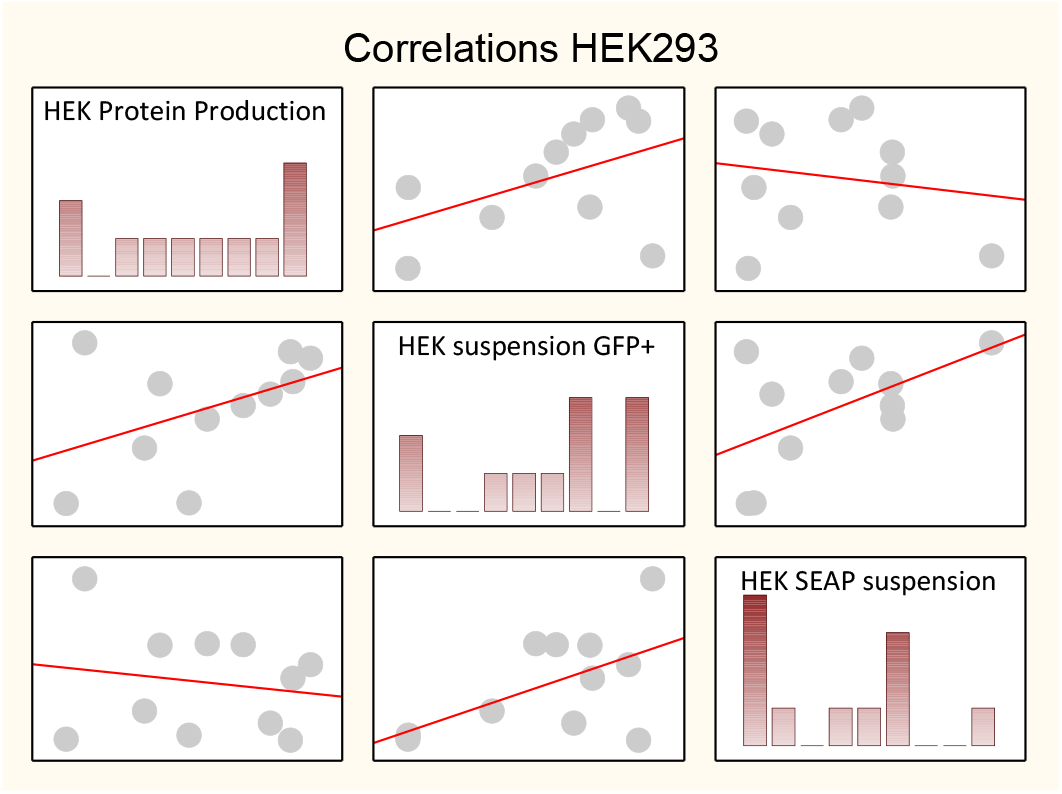
Scatterplot that highlights the correlations between the protein production yields and the assays of transfection-positive cell population and SEAP protein secretion in HEK293 cells.

