## Supplementary Table 2 for "Expressed therapeutic protein yields are predicted by transiently transfected mammalian cell population": Supplementary_Table_2.pdf

#### Statistical analysis of the transfection efficacy

##### Total reporter from the cell lysate in CHO adherent culture.

Statistical analysis of the variance (1-way ANOVA) for the dataset presented in Figure 1A is presented below, together with Tukey post-hoc test p values.

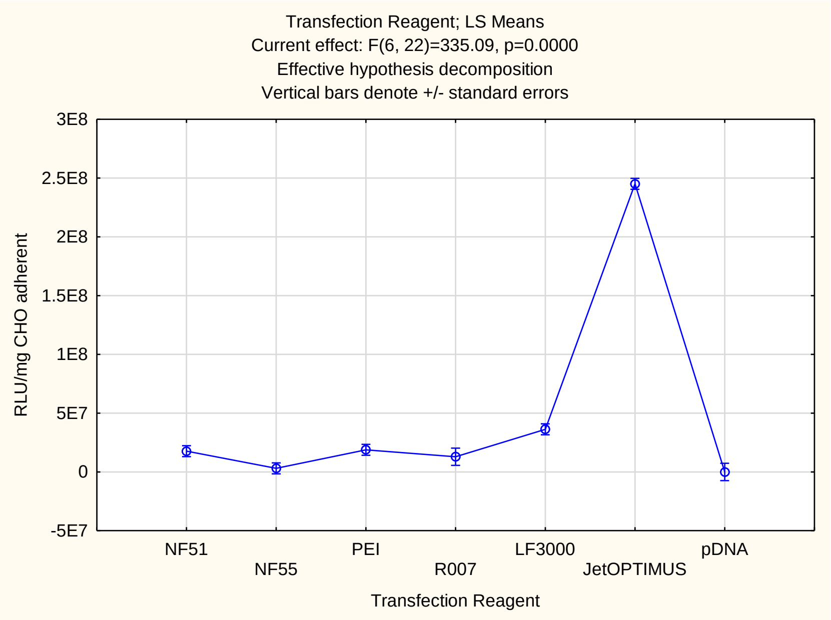


Tukey HSD test; variable RLU/mg CHO adherent(Spreadsheet12 in 20220215 Fig1A statistics)
Approximate Probabilities for Post Hoc Tests
Error: Between MS = 109E12, df = 22.000

| Cell No. | Transfection Reagent | {1} 1766E4 | {2} 3050E3 | {3} 1878E4 | {4} 1291E4 | {5} 3626E4 | {6} 2452E5 | {7} 1715.0 |
|---|---|---|---|---|---|---|---|---|
| 1 | NF51 | | 0.329 | 1.000 | 0.998 | 0.116 | 0.000 | 0.430 |
| 2 | NF55 | 0.329 | | 0.252 | 0.912 | 0.001 | 0.000 | 1.000 |
| 3 | PEI | 1.000 | 0.252 | | 0.993 | 0.160 | 0.000 | 0.360 |
| 4 | R007 | 0.998 | 0.912 | 0.993 | | 0.153 | 0.000 | 0.872 |
| 5 | LF3000 | 0.116 | 0.001 | 0.160 | 0.153 | | 0.000 | 0.007 |
| 6 | JetOPTIMUS | 0.000 | 0.000 | 0.000 | 0.000 | 0.000 | | 0.000 |
| 7 | pDNA | 0.430 | 1.000 | 0.360 | 0.872 | 0.007 | 0.000 | |

#### Statistical analysis of the transfection efficacy

##### Total reporter from the cell lysate in CHO adherent culture. Extended set of transfection reagents.

Statistical analysis of the variance (1-way ANOVA) for the dataset presented in Figure 2A is presented below, together with Tukey post-hoc test p values.

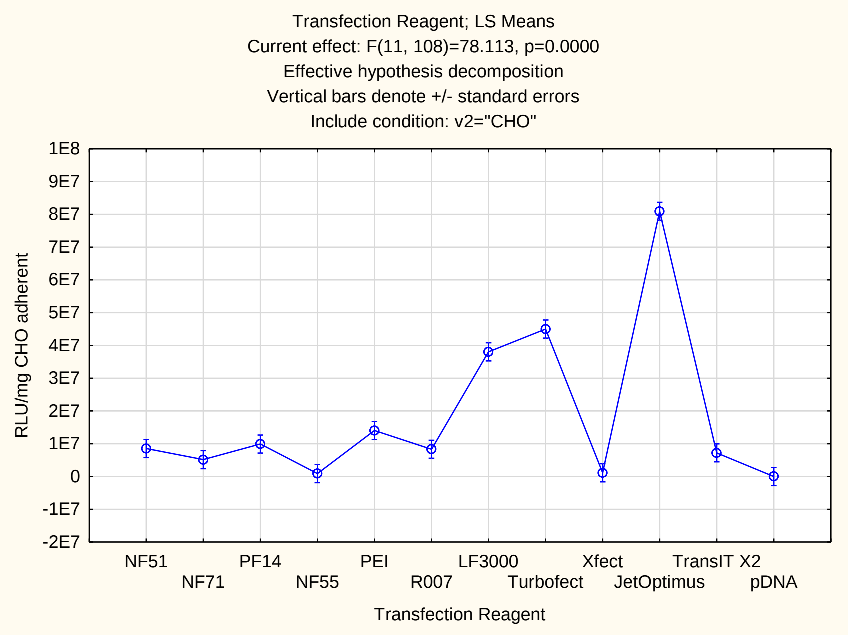

Tukey HSD test; variable RLU/mg CHO adherent
Approximate Probabilities for Post Hoc Tests
Error: Between MS = 761E11, df = 108.00
Include condition: v2="CHO"

| Cell No. | Transfection Reagent | {1} 8528E3 | {2} 5153E3 | {3} 9915E3 | {4} 9014E2 | {5} 1403E4 | {6} 8316E3 | {7} 3805E4 | {8} 4501E4 | {9} 1138E3 | {10} 8095E4 | {11} 7215E3 | {12} 239.93 |
|---|---|---|---|---|---|---|---|---|---|---|---|---|---|
| 1 | NF51 | | 0.999 | 1.000 | 0.722 | 0.959 | 1.000 | 0.000 | 0.000 | 0.760 | 0.000 | 1.000 | 0.564 |
| 2 | NF71 | 0.999 | | 0.986 | 0.995 | 0.500 | 1.000 | 0.000 | 0.000 | 0.997 | 0.000 | 1.000 | 0.975 |
| 3 | PF14 | 1.000 | 0.986 | | 0.476 | 0.996 | 1.000 | 0.000 | 0.000 | 0.518 | 0.000 | 1.000 | 0.327 |
| 4 | NF55 | 0.722 | 0.995 | 0.476 | | 0.047 | 0.757 | 0.000 | 0.000 | 1.000 | 0.000 | 0.899 | 1.000 |
| 5 | PEI | 0.959 | 0.500 | 0.996 | 0.047 | | 0.947 | 0.000 | 0.000 | 0.055 | 0.000 | 0.842 | 0.024 |
| 6 | R007 | 1.000 | 1.000 | 1.000 | 0.757 | 0.947 | | 0.000 | 0.000 | 0.792 | 0.000 | 1.000 | 0.602 |
| 7 | LF3000 | 0.000 | 0.000 | 0.000 | 0.000 | 0.000 | 0.000 | | 0.824 | 0.000 | 0.000 | 0.000 | 0.000 |
| 8 | Turbofect | 0.000 | 0.000 | 0.000 | 0.000 | 0.000 | 0.000 | 0.824 | | 0.000 | 0.000 | 0.000 | 0.000 |
| 9 | Xfect | 0.760 | 0.997 | 0.518 | 1.000 | 0.055 | 0.792 | 0.000 | 0.000 | | 0.000 | 0.920 | 1.000 |
| 10 | JetOptimus | 0.000 | 0.000 | 0.000 | 0.000 | 0.000 | 0.000 | 0.000 | 0.000 | 0.000 | | 0.000 | 0.000 |
| 11 | TransIT X2 | 1.000 | 1.000 | 1.000 | 0.899 | 0.842 | 1.000 | 0.000 | 0.000 | 0.920 | 0.000 | | 0.787 |
| 12 | pDNA | 0.564 | 0.975 | 0.327 | 1.000 | 0.024 | 0.602 | 0.000 | 0.000 | 1.000 | 0.000 | 0.787 | |

### Statistical analysis of the transfection efficacy

#### Total reporter from the cell lysate in HEK293 adherent culture.

Statistical analysis of the variance (1-way ANOVA) for the dataset presented in Figure 2A is presented below, together with Tukey post-hoc test p values.

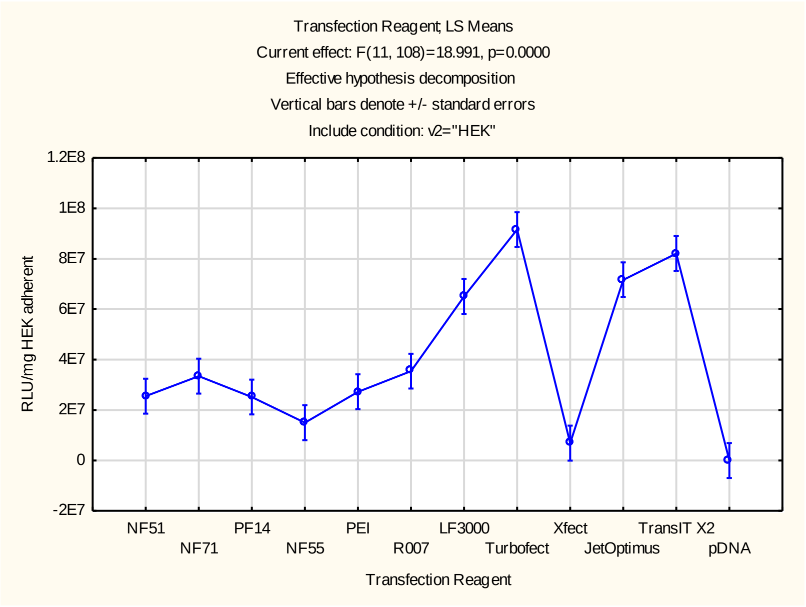


Tukey HSD test; variable RLU/mg CHO adherent
Approximate Probabilities for Post Hoc Tests
Error: Between MS = 480E12, df = 108.00
Include condition: v2="HEK"

| Cell No. | Transfection Reagent | {1} 2552E4 | {2} 3347E4 | {3} 2517E4 | {4} 1497E4 | {5} 2723E4 | {6} 3543E4 | {7} 6505E4 | {8} 9157E4 | {9} 6876E3 | {10} 7166E4 | {11} 8205E4 | {12} 306.80 |
|---|---|---|---|---|---|---|---|---|---|---|---|---|---|
| 1 | NF51 | | 1.000 | 1.000 | 0.995 | 1.000 | 0.997 | 0.006 | 0.000 | 0.756 | 0.001 | 0.000 | 0.291 |
| 2 | NF71 | 1.000 | | 0.999 | 0.764 | 1.000 | 1.000 | 0.070 | 0.000 | 0.234 | 0.009 | 0.000 | 0.041 |
| 3 | PF14 | 1.000 | 0.999 | | 0.996 | 1.000 | 0.996 | 0.005 | 0.000 | 0.777 | 0.001 | 0.000 | 0.312 |
| 4 | NF55 | 0.995 | 0.764 | 0.996 | | 0.983 | 0.633 | 0.000 | 0.000 | 1.000 | 0.000 | 0.000 | 0.930 |
| 5 | PEI | 1.000 | 1.000 | 1.000 | 0.983 | | 1.000 | 0.010 | 0.000 | 0.641 | 0.001 | 0.000 | 0.205 |
| 6 | R007 | 0.997 | 1.000 | 0.996 | 0.633 | 1.000 | | 0.117 | 0.000 | 0.151 | 0.017 | 0.000 | 0.022 |
| 7 | LF3000 | 0.006 | 0.070 | 0.005 | 0.000 | 0.010 | 0.117 | | 0.238 | 0.000 | 1.000 | 0.848 | 0.000 |
| 8 | Turbofect | 0.000 | 0.000 | 0.000 | 0.000 | 0.000 | 0.000 | 0.238 | | 0.000 | 0.671 | 0.998 | 0.000 |
| 9 | Xfect | 0.756 | 0.234 | 0.777 | 1.000 | 0.641 | 0.151 | 0.000 | 0.000 | | 0.000 | 0.000 | 1.000 |
| 10 | JetOptimus | 0.001 | 0.009 | 0.001 | 0.000 | 0.001 | 0.017 | 1.000 | 0.671 | 0.000 | | 0.996 | 0.000 |
| 11 | TransIT X2 | 0.000 | 0.000 | 0.000 | 0.000 | 0.000 | 0.000 | 0.848 | 0.998 | 0.000 | 0.996 | | 0.000 |
| 12 | pDNA | 0.291 | 0.041 | 0.312 | 0.930 | 0.205 | 0.022 | 0.000 | 0.000 | 1.000 | 0.000 | 0.000 | |

#### Statistical analysis of the transfection efficacy

##### Total reporter from the cell lysate in CHO suspension culture.

Statistical analysis of the variance (1-way ANOVA) for the dataset presented in Figure 2B is presented below, together with Tukey post-hoc test p values.

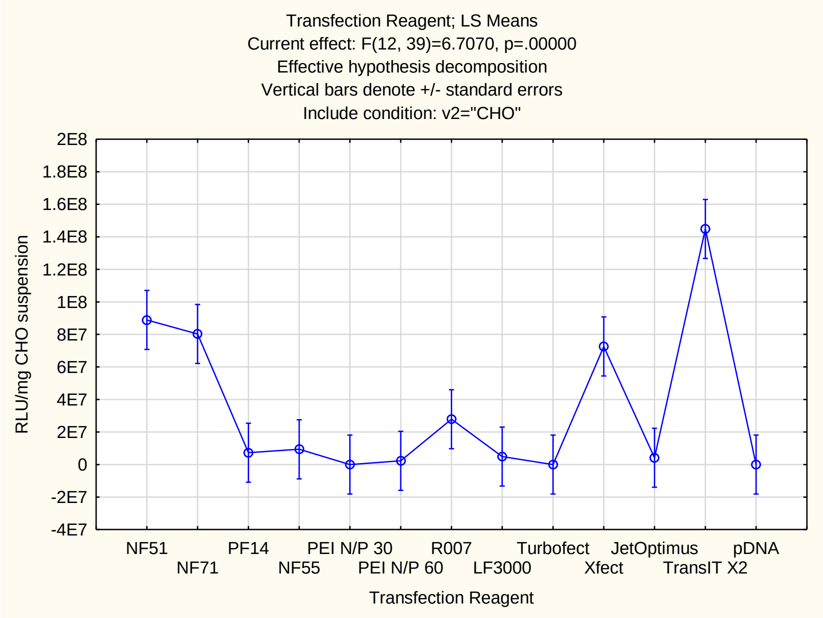

| | Tukey HSD test; variable RLU/mg suspension<br>Approximate Probabilities for Post Hoc Tests<br>Error: Between MS = 132E13, df = 39.000<br>Include condition: v2="CHO" | | | | | | | | | | | | | |
|---|---|---|---|---|---|---|---|---|---|---|---|---|---|---|
| Cell No. | Transfection Reagent | {1}<br>8890E4 | {2}<br>8029E4 | {3}<br>7285E3 | {4}<br>9392E3 | {5}<br>3411.7 | {6}<br>2304E3 | {7}<br>2788E4 | {8}<br>4898E3 | {9}<br>2644.1 | {10}<br>7261E4 | {11}<br>4183E3 | {12}<br>1448E5 | {13}<br>1038.3 |
| 1 | NF51 | | 1.000 | 0.112 | 0.134 | 0.058 | 0.072 | 0.479 | 0.091 | 0.058 | 1.000 | 0.085 | 0.610 | 0.058 |
| 2 | NF71 | 1.000 | | 0.223 | 0.260 | 0.125 | 0.152 | 0.700 | 0.186 | 0.125 | 1.000 | 0.176 | 0.393 | 0.125 |
| 3 | PF14 | 0.112 | 0.223 | | 1.000 | 1.000 | 1.000 | 1.000 | 1.000 | 1.000 | 0.375 | 1.000 | 0.000 | 1.000 |
| 4 | NF55 | 0.134 | 0.260 | 1.000 | | 1.000 | 1.000 | 1.000 | 1.000 | 1.000 | 0.425 | 1.000 | 0.000 | 1.000 |
| 5 | PEI N/P 30 | 0.058 | 0.125 | 1.000 | 1.000 | | 1.000 | 0.996 | 1.000 | 1.000 | 0.230 | 1.000 | 0.000 | 1.000 |
| 6 | PEI N/P 60 | 0.072 | 0.152 | 1.000 | 1.000 | 1.000 | | 0.998 | 1.000 | 1.000 | 0.271 | 1.000 | 0.000 | 1.000 |
| 7 | R007 | 0.479 | 0.700 | 1.000 | 1.000 | 0.996 | 0.998 | | 0.999 | 0.996 | 0.865 | 0.999 | 0.003 | 0.996 |
| 8 | LF3000 | 0.091 | 0.186 | 1.000 | 1.000 | 1.000 | 1.000 | 0.999 | | 1.000 | 0.323 | 1.000 | 0.000 | 1.000 |
| 9 | Turbofect | 0.058 | 0.125 | 1.000 | 1.000 | 1.000 | 1.000 | 0.996 | 1.000 | | 0.230 | 1.000 | 0.000 | 1.000 |
| 10 | Xfect | 1.000 | 1.000 | 0.375 | 0.425 | 0.230 | 0.271 | 0.865 | 0.323 | 0.230 | | 0.308 | 0.236 | 0.230 |
| 11 | JetOptimus | 0.085 | 0.176 | 1.000 | 1.000 | 1.000 | 1.000 | 0.999 | 1.000 | 1.000 | 0.308 | | 0.000 | 1.000 |
| 12 | TransIT X2 | 0.610 | 0.393 | 0.000 | 0.000 | 0.000 | 0.000 | 0.003 | 0.000 | 0.000 | 0.236 | 0.000 | | 0.000 |
| 13 | pDNA | 0.058 | 0.125 | 1.000 | 1.000 | 1.000 | 1.000 | 0.996 | 1.000 | 1.000 | 0.230 | 1.000 | 0.000 | |

#### Statistical analysis of the transfection efficacy

##### Total reporter from the cell lysate in HEK293 suspension culture.

Statistical analysis of the variance (1-way ANOVA) for the dataset presented in Figure 2B is presented below, together with Tukey post-hoc test p values.

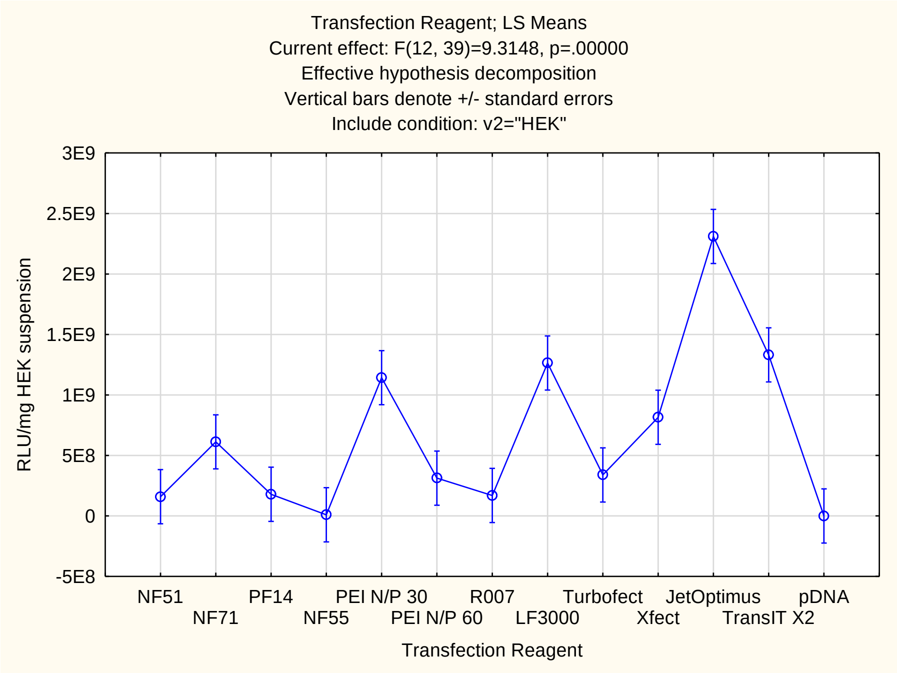


Tukey HSD test; variable RLU/mg suspension
Approximate Probabilities for Post Hoc Tests
Error: Between MS = 201E15, df = 39.000
Include condition: v2="HEK"

| Cell No. | Transfection Reagent | {1} 1594E5 | {2} 6124E5 | {3} 1793E5 | {4} 9611E3 | {5} 1143E6 | {6} 3125E5 | {7} 1691E5 | {8} 1264E6 | {9} 3392E5 | {10} 8156E5 | {11} 2310E6 | {12} 1332E6 | {13} 25668. |
|---|---|---|---|---|---|---|---|---|---|---|---|---|---|---|
| 1 | NF51 | | 0.963 | 1.000 | 1.000 | 0.132 | 1.000 | 1.000 | 0.055 | 1.000 | 0.681 | 0.000 | 0.032 | 1.000 |
| 2 | NF71 | 0.963 | | 0.973 | 0.783 | 0.892 | 0.999 | 0.968 | 0.690 | 1.000 | 1.000 | 0.000 | 0.550 | 0.766 |
| 3 | PF14 | 1.000 | 0.973 | | 1.000 | 0.150 | 1.000 | 1.000 | 0.064 | 1.000 | 0.721 | 0.000 | 0.038 | 1.000 |
| 4 | NF55 | 1.000 | 0.783 | 1.000 | | 0.044 | 0.999 | 1.000 | 0.016 | 0.997 | 0.375 | 0.000 | 0.009 | 1.000 |
| 5 | PEI N/P 30 | 0.132 | 0.892 | 0.150 | 0.044 | | 0.331 | 0.140 | 1.000 | 0.379 | 0.997 | 0.033 | 1.000 | 0.040 |
| 6 | PEI N/P 60 | 1.000 | 0.999 | 1.000 | 0.999 | 0.331 | | 1.000 | 0.163 | 1.000 | 0.923 | 0.000 | 0.103 | 0.998 |
| 7 | R007 | 1.000 | 0.968 | 1.000 | 1.000 | 0.140 | 1.000 | | 0.059 | 1.000 | 0.700 | 0.000 | 0.035 | 1.000 |
| 8 | LF3000 | 0.055 | 0.690 | 0.064 | 0.016 | 1.000 | 0.163 | 0.059 | | 0.193 | 0.965 | 0.085 | 1.000 | 0.015 |
| 9 | Turbofect | 1.000 | 1.000 | 1.000 | 0.997 | 0.379 | 1.000 | 1.000 | 0.193 | | 0.946 | 0.000 | 0.124 | 0.997 |
| 10 | Xfect | 0.681 | 1.000 | 0.721 | 0.375 | 0.997 | 0.923 | 0.700 | 0.965 | 0.946 | | 0.002 | 0.910 | 0.358 |
| 11 | JetOptimus | 0.000 | 0.000 | 0.000 | 0.000 | 0.033 | 0.000 | 0.000 | 0.085 | 0.000 | 0.002 | | 0.136 | 0.000 |
| 12 | TransIT X2 | 0.032 | 0.550 | 0.038 | 0.009 | 1.000 | 0.103 | 0.035 | 1.000 | 0.124 | 0.910 | 0.136 | | 0.008 |
| 13 | pDNA | 1.000 | 0.766 | 1.000 | 1.000 | 0.040 | 0.998 | 1.000 | 0.015 | 0.997 | 0.358 | 0.000 | 0.008 | |

#### Statistical analysis of the transfection efficacy

##### Transfection-positive cell population in CHO adherent culture.

Statistical analysis of the variance (1-way ANOVA) for the dataset presented in Figure 2C is presented below, together with Tukey post-hoc test p values.

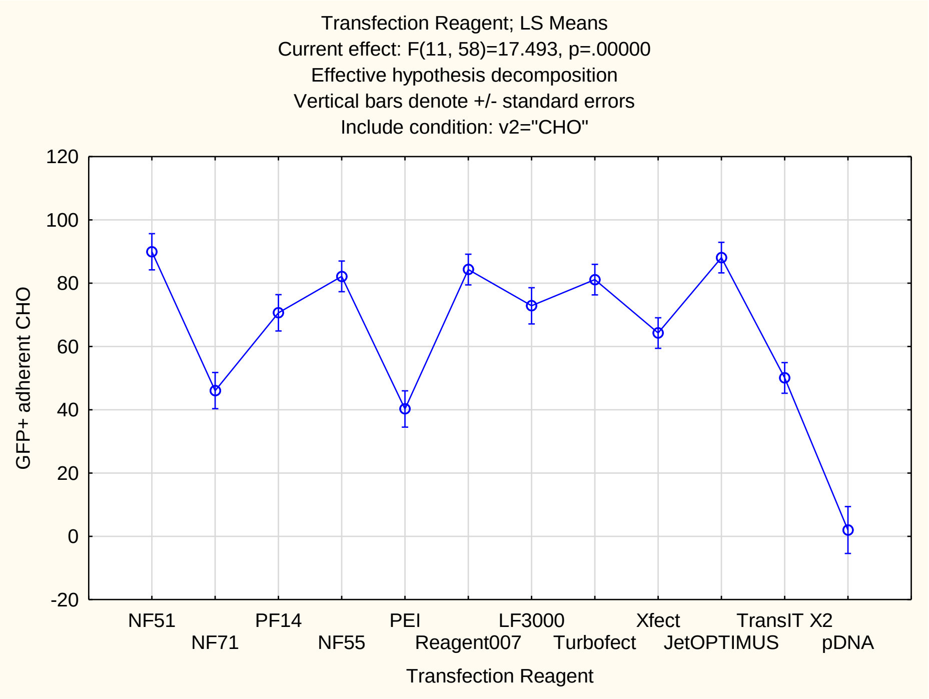

Tukey HSD test; variable GFP+ adherent
Approximate Probabilities for Post Hoc Tests
Error: Between MS = 163.95, df = 58.000
Include condition: v2="CHO"

| Cell No. | Transfection Reagent | {1} 89.958 | {2} 46.046 | {3} 70.645 | {4} 82.163 | {5} 40.245 | {6} 84.318 | {7} 72.867 | {8} 81.137 | {9} 64.249 | {10} 88.089 | {11} 50.068 | {12} 1.9890 |
|---|---|---|---|---|---|---|---|---|---|---|---|---|---|
| 1 | NF51 | | 0.000 | 0.432 | 0.996 | 0.000 | 1.000 | 0.618 | 0.989 | 0.047 | 1.000 | 0.000 | 0.000 |
| 2 | NF71 | 0.000 | | 0.124 | 0.001 | 1.000 | 0.000 | 0.064 | 0.001 | 0.405 | 0.000 | 1.000 | 0.001 |
| 3 | PF14 | 0.432 | 0.124 | | 0.924 | 0.019 | 0.798 | 1.000 | 0.959 | 0.999 | 0.471 | 0.232 | 0.000 |
| 4 | NF55 | 0.996 | 0.001 | 0.924 | | 0.000 | 1.000 | 0.983 | 1.000 | 0.294 | 0.999 | 0.001 | 0.000 |
| 5 | PEI | 0.000 | 1.000 | 0.019 | 0.000 | | 0.000 | 0.008 | 0.000 | 0.084 | 0.000 | 0.974 | 0.007 |
| 6 | Reagent007 | 1.000 | 0.000 | 0.798 | 1.000 | 0.000 | | 0.927 | 1.000 | 0.157 | 1.000 | 0.000 | 0.000 |
| 7 | LF3000 | 0.618 | 0.064 | 1.000 | 0.983 | 0.008 | 0.927 | | 0.993 | 0.991 | 0.672 | 0.123 | 0.000 |
| 8 | Turbofect | 0.989 | 0.001 | 0.959 | 1.000 | 0.000 | 1.000 | 0.993 | | 0.380 | 0.997 | 0.002 | 0.000 |
| 9 | Xfect | 0.047 | 0.405 | 0.999 | 0.294 | 0.084 | 0.157 | 0.991 | 0.380 | | 0.041 | 0.644 | 0.000 |
| 10 | JetOPTIMUS | 1.000 | 0.000 | 0.471 | 0.999 | 0.000 | 1.000 | 0.672 | 0.997 | 0.041 | | 0.000 | 0.000 |
| 11 | TransIT X2 | 0.000 | 1.000 | 0.232 | 0.001 | 0.974 | 0.000 | 0.123 | 0.002 | 0.644 | 0.000 | | 0.000 |
| 12 | pDNA | 0.000 | 0.001 | 0.000 | 0.000 | 0.007 | 0.000 | 0.000 | 0.000 | 0.000 | 0.000 | 0.000 | |

### Statistical analysis of the transfection efficacy

#### Transfection-positive cell population in HEK293 adherent culture.

Statistical analysis of the variance (1-way ANOVA) for the dataset presented in Figure 2C is presented below, together with Tukey post-hoc test p values.

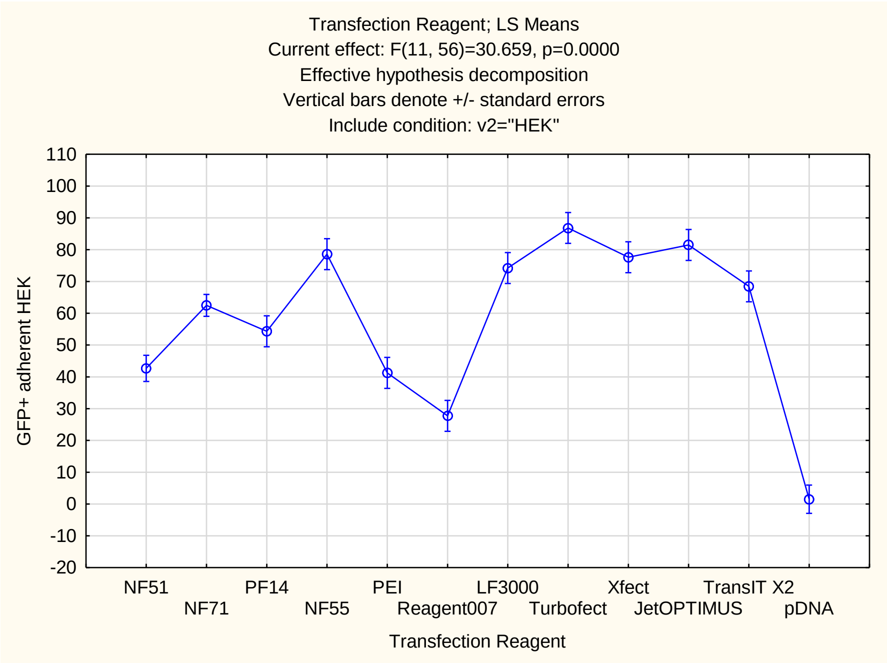


Tukey HSD test; variable GFP+ adherent
Approximate Probabilities for Post Hoc Tests
Error: Between MS = 118.10, df = 56.000
Include condition: v2="HEK"

| Cell No. | Transfection Reagent | {1} 42.670 | {2} 62.481 | {3} 54.338 | {4} 78.594 | {5} 41.244 | {6} 27.726 | {7} 74.236 | {8} 86.836 | {9} 77.639 | {10} 81.480 | {11} 68.457 | {12} 1.5035 |
|---|---|---|---|---|---|---|---|---|---|---|---|---|---|
| 1 | NF51 | | 0.023 | 0.793 | 0.000 | 1.000 | 0.457 | 0.001 | 0.000 | 0.000 | 0.000 | 0.008 | 0.000 |
| 2 | NF71 | 0.023 | | 0.965 | 0.250 | 0.033 | 0.000 | 0.708 | 0.007 | 0.334 | 0.087 | 0.997 | 0.000 |
| 3 | PF14 | 0.793 | 0.965 | | 0.036 | 0.751 | 0.014 | 0.171 | 0.001 | 0.053 | 0.011 | 0.656 | 0.000 |
| 4 | NF55 | 0.000 | 0.250 | 0.036 | | 0.000 | 0.000 | 1.000 | 0.987 | 1.000 | 1.000 | 0.941 | 0.000 |
| 5 | PEI | 1.000 | 0.033 | 0.751 | 0.000 | | 0.713 | 0.001 | 0.000 | 0.000 | 0.000 | 0.011 | 0.000 |
| 6 | Reagent007 | 0.457 | 0.000 | 0.014 | 0.000 | 0.713 | | 0.000 | 0.000 | 0.000 | 0.000 | 0.000 | 0.010 |
| 7 | LF3000 | 0.001 | 0.708 | 0.171 | 1.000 | 0.001 | 0.000 | | 0.793 | 1.000 | 0.995 | 0.999 | 0.000 |
| 8 | Turbofect | 0.000 | 0.007 | 0.001 | 0.987 | 0.000 | 0.000 | 0.793 | | 0.970 | 1.000 | 0.266 | 0.000 |
| 9 | Xfect | 0.000 | 0.334 | 0.053 | 1.000 | 0.000 | 0.000 | 1.000 | 0.970 | | 1.000 | 0.970 | 0.000 |
| 10 | JetOPTIMUS | 0.000 | 0.087 | 0.011 | 1.000 | 0.000 | 0.000 | 0.995 | 1.000 | 1.000 | | 0.757 | 0.000 |
| 11 | TransIT X2 | 0.008 | 0.997 | 0.656 | 0.941 | 0.011 | 0.000 | 0.999 | 0.266 | 0.970 | 0.757 | | 0.000 |
| 12 | pDNA | 0.000 | 0.000 | 0.000 | 0.000 | 0.000 | 0.010 | 0.000 | 0.000 | 0.000 | 0.000 | 0.000 | |

#### Statistical analysis of the transfection efficacy

##### Transfection-positive cell population in CHO suspension culture.

Statistical analysis of the variance (1-way ANOVA) for the dataset presented in Figure 2D is presented below, together with Tukey post-hoc test p values.

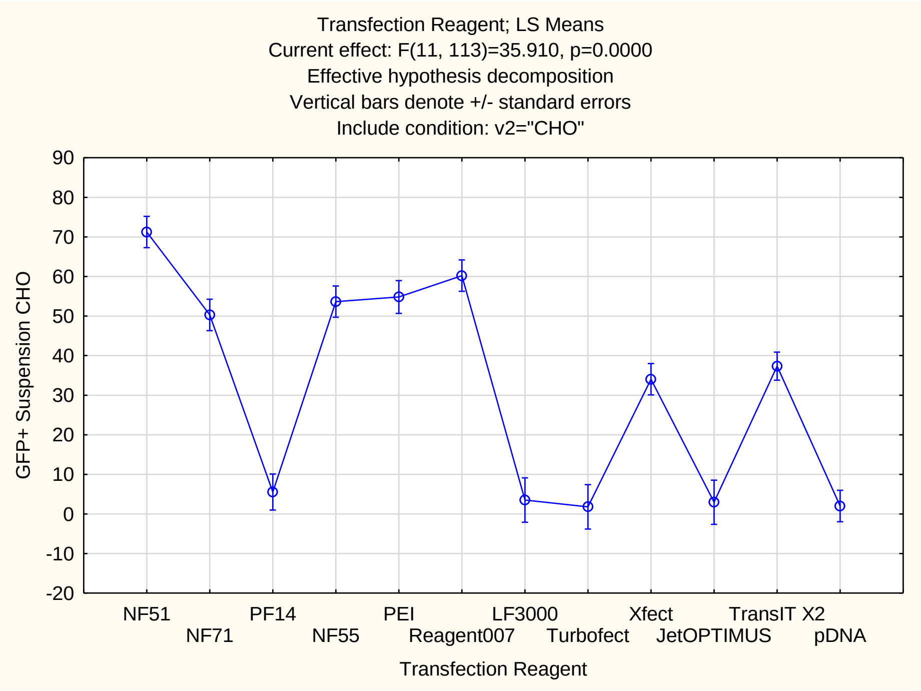

Tukey HSD test; variable GFP+ Suspension
Approximate Probabilities for Post Hoc Tests
Error: Between MS = 187.87, df = 113.00
Include condition: v2="CHO"

| Cell No. | Transfection Reagent | {1} 71.247 | {2} 50.300 | {3} 5.5441 | {4} 53.650 | {5} 54.825 | {6} 60.216 | {7} 3.5202 | {8} 1.7970 | {9} 34.047 | {10} 2.9652 | {11} 37.342 | {12} 2.0138 |
|---|---|---|---|---|---|---|---|---|---|---|---|---|---|
| 1 | NF51 | | 0.015 | 0.000 | 0.085 | 0.166 | 0.711 | 0.000 | 0.000 | 0.000 | 0.000 | 0.000 | 0.000 |
| 2 | NF71 | 0.015 | | 0.000 | 1.000 | 1.000 | 0.829 | 0.000 | 0.000 | 0.154 | 0.000 | 0.388 | 0.000 |
| 3 | PF14 | 0.000 | 0.000 | | 0.000 | 0.000 | 0.000 | 1.000 | 1.000 | 0.001 | 1.000 | 0.000 | 1.000 |
| 4 | NF55 | 0.085 | 1.000 | 0.000 | | 1.000 | 0.990 | 0.000 | 0.000 | 0.031 | 0.000 | 0.102 | 0.000 |
| 5 | PEI | 0.166 | 1.000 | 0.000 | 1.000 | | 0.999 | 0.000 | 0.000 | 0.021 | 0.000 | 0.071 | 0.000 |
| 6 | Reagent007 | 0.711 | 0.829 | 0.000 | 0.990 | 0.999 | | 0.000 | 0.000 | 0.001 | 0.000 | 0.002 | 0.000 |
| 7 | LF3000 | 0.000 | 0.000 | 1.000 | 0.000 | 0.000 | 0.000 | | 1.000 | 0.001 | 1.000 | 0.000 | 1.000 |
| 8 | Turbofect | 0.000 | 0.000 | 1.000 | 0.000 | 0.000 | 0.000 | 1.000 | | 0.001 | 1.000 | 0.000 | 1.000 |
| 9 | Xfect | 0.000 | 0.154 | 0.001 | 0.031 | 0.021 | 0.001 | 0.001 | 0.001 | | 0.001 | 1.000 | 0.000 |
| 10 | JetOPTIMUS | 0.000 | 0.000 | 1.000 | 0.000 | 0.000 | 0.000 | 1.000 | 1.000 | 0.001 | | 0.000 | 1.000 |
| 11 | TransIT X2 | 0.000 | 0.388 | 0.000 | 0.102 | 0.071 | 0.002 | 0.000 | 0.000 | 1.000 | 0.000 | | 0.000 |
| 12 | pDNA | 0.000 | 0.000 | 1.000 | 0.000 | 0.000 | 0.000 | 1.000 | 1.000 | 0.000 | 1.000 | 0.000 | |

#### Statistical analysis of the transfection efficacy

##### Transfection-positive cell population in HEK293 suspension culture.

Statistical analysis of the variance (1-way ANOVA) for the dataset presented in Figure 2D is presented below, together with Tukey post-hoc test p values.

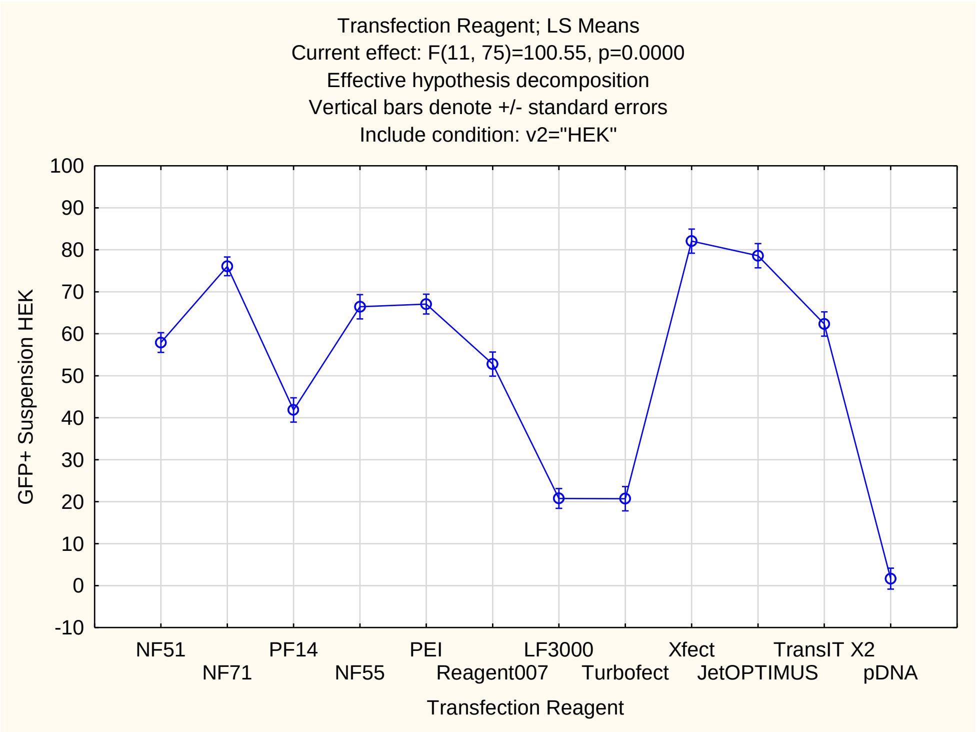


Tukey HSD test; variable GFP+ Suspension
Approximate Probabilities for Post Hoc Tests
Error: Between MS = 49.947, df = 75.000
Include condition: v2="HEK"

| Cell No. | Transfection Reagent | {1} 57.922 | {2} 76.076 | {3} 41.856 | {4} 66.428 | {5} 67.071 | {6} 52.777 | {7} 20.779 | {8} 20.710 | {9} 82.072 | {10} 78.592 | {11} 62.335 | {12} 1.6703 |
|---|---|---|---|---|---|---|---|---|---|---|---|---|---|
| 1 | NF51 | | 0.000 | 0.003 | 0.497 | 0.225 | 0.964 | 0.000 | 0.000 | 0.000 | 0.000 | 0.989 | 0.000 |
| 2 | NF71 | 0.000 | | 0.000 | 0.275 | 0.213 | 0.000 | 0.000 | 0.000 | 0.887 | 1.000 | 0.016 | 0.000 |
| 3 | PF14 | 0.003 | 0.000 | | 0.000 | 0.000 | 0.259 | 0.000 | 0.000 | 0.000 | 0.000 | 0.000 | 0.000 |
| 4 | NF55 | 0.497 | 0.275 | 0.000 | | 1.000 | 0.054 | 0.000 | 0.000 | 0.013 | 0.135 | 0.997 | 0.000 |
| 5 | PEI | 0.225 | 0.213 | 0.000 | 1.000 | | 0.013 | 0.000 | 0.000 | 0.007 | 0.103 | 0.980 | 0.000 |
| 6 | Reagent007 | 0.964 | 0.000 | 0.259 | 0.054 | 0.013 | | 0.000 | 0.000 | 0.000 | 0.000 | 0.457 | 0.000 |
| 7 | LF3000 | 0.000 | 0.000 | 0.000 | 0.000 | 0.000 | 0.000 | | 1.000 | 0.000 | 0.000 | 0.000 | 0.000 |
| 8 | Turbofect | 0.000 | 0.000 | 0.000 | 0.000 | 0.000 | 0.000 | 1.000 | | 0.000 | 0.000 | 0.000 | 0.000 |
| 9 | Xfect | 0.000 | 0.887 | 0.000 | 0.013 | 0.007 | 0.000 | 0.000 | 0.000 | | 0.999 | 0.001 | 0.000 |
| 10 | JetOPTIMUS | 0.000 | 1.000 | 0.000 | 0.135 | 0.103 | 0.000 | 0.000 | 0.000 | 0.999 | | 0.008 | 0.000 |
| 11 | TransIT X2 | 0.989 | 0.016 | 0.000 | 0.997 | 0.980 | 0.457 | 0.000 | 0.000 | 0.001 | 0.008 | | 0.000 |
| 12 | pDNA | 0.000 | 0.000 | 0.000 | 0.000 | 0.000 | 0.000 | 0.000 | 0.000 | 0.000 | 0.000 | 0.000 | |

#### Statistical analysis of the transfection efficacy

##### Total secreted protein in CHO suspension culture.

Statistical analysis of the variance (1-way ANOVA) for the dataset presented in Figure 3A is presented below, together with Tukey post-hoc test p values.

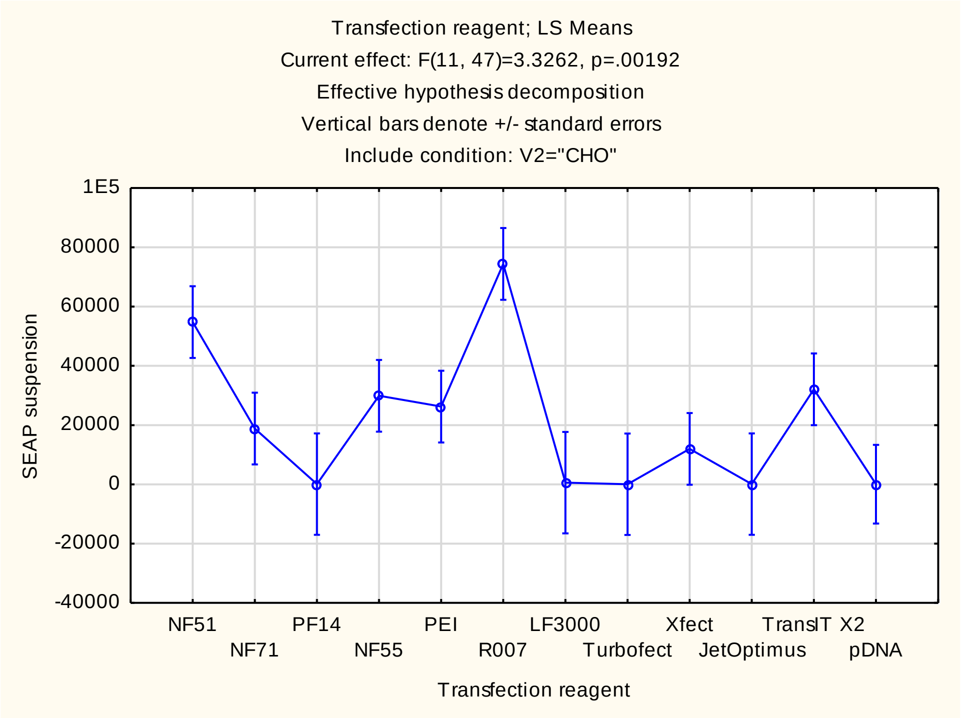

Tukey HSD test; variable SEAP suspension
Approximate Probabilities for Post Hoc Tests
Error: Between MS = 8783E5, df = 47.000
Include condition: V2="CHO"

| Cell No. | Transfection reagent | {1} 54775. | {2} 18828. | {3} 104.00 | {4} 29900. | {5} 26256. | {6} 74385. | {7} 591.00 | {8} 48.333 | {9} 11978. | {10} 68.667 | {11} 32092. | {12} 67.200 |
|---|---|---|---|---|---|---|---|---|---|---|---|---|---|
| 1 | NF51 | | 0.625 | 0.304 | 0.946 | 0.874 | 0.991 | 0.317 | 0.303 | 0.364 | 0.303 | 0.971 | 0.127 |
| 2 | NF71 | 0.625 | | 0.999 | 1.000 | 1.000 | 0.080 | 0.999 | 0.999 | 1.000 | 0.999 | 1.000 | 0.996 |
| 3 | PF14 | 0.304 | 0.999 | | 0.953 | 0.982 | 0.038 | 1.000 | 1.000 | 1.000 | 1.000 | 0.925 | 1.000 |
| 4 | NF55 | 0.946 | 1.000 | 0.953 | | 1.000 | 0.309 | 0.958 | 0.953 | 0.996 | 0.953 | 1.000 | 0.876 |
| 5 | PEI | 0.874 | 1.000 | 0.982 | 1.000 | | 0.208 | 0.984 | 0.981 | 0.999 | 0.981 | 1.000 | 0.944 |
| 6 | R007 | 0.991 | 0.080 | 0.038 | 0.309 | 0.208 | | 0.040 | 0.038 | 0.029 | 0.038 | 0.382 | 0.007 |
| 7 | LF3000 | 0.317 | 0.999 | 1.000 | 0.958 | 0.984 | 0.040 | | 1.000 | 1.000 | 1.000 | 0.932 | 1.000 |
| 8 | Turbofec | 0.303 | 0.999 | 1.000 | 0.953 | 0.981 | 0.038 | 1.000 | | 1.000 | 1.000 | 0.925 | 1.000 |
| 9 | Xfect | 0.364 | 1.000 | 1.000 | 0.996 | 0.999 | 0.029 | 1.000 | 1.000 | | 1.000 | 0.988 | 1.000 |
| 10 | JetOptimu | 0.303 | 0.999 | 1.000 | 0.953 | 0.981 | 0.038 | 1.000 | 1.000 | 1.000 | | 0.925 | 1.000 |
| 11 | TransIT X | 0.971 | 1.000 | 0.925 | 1.000 | 1.000 | 0.382 | 0.932 | 0.925 | 0.988 | 0.925 | | 0.818 |
| 12 | pDNA | 0.127 | 0.996 | 1.000 | 0.876 | 0.944 | 0.007 | 1.000 | 1.000 | 1.000 | 1.000 | 0.818 | |

#### Statistical analysis of the transfection efficacy

##### Total secreted protein in HEK293 suspension culture.

Statistical analysis of the variance (1-way ANOVA) for the dataset presented in Figure 3A is presented below, together with Tukey post-hoc test p values.

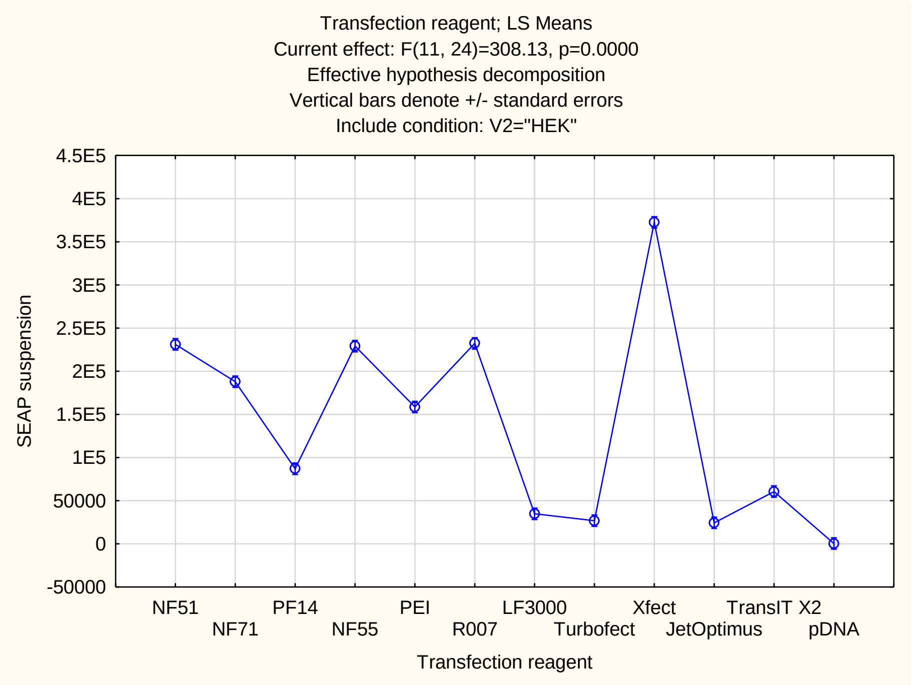


Tukey HSD test; variable SEAP suspension
Approximate Probabilities for Post Hoc Tests
Error: Between MS = 1302E5, df = 24.000
Include condition: V2="HEK"

| Cell No. | Transfection reagent | {1} 2311E | {2} 1879E | {3} 87038. | {4} 2291E | {5} 1586E | {6} 2323E | {7} 34843. | {8} 26884. | {9} 3725E | {10} 24405. | {11} 60608. | {12} 433.33 |
|---|---|---|---|---|---|---|---|---|---|---|---|---|---|
| 1 | NF51 | | 0.005 | 0.000 | 1.000 | 0.000 | 1.000 | 0.000 | 0.000 | 0.000 | 0.000 | 0.000 | 0.000 |
| 2 | NF71 | 0.005 | | 0.000 | 0.008 | 0.131 | 0.004 | 0.000 | 0.000 | 0.000 | 0.000 | 0.000 | 0.000 |
| 3 | PF14 | 0.000 | 0.000 | | 0.000 | 0.000 | 0.000 | 0.001 | 0.000 | 0.000 | 0.000 | 0.226 | 0.000 |
| 4 | NF55 | 1.000 | 0.008 | 0.000 | | 0.000 | 1.000 | 0.000 | 0.000 | 0.000 | 0.000 | 0.000 | 0.000 |
| 5 | PEI | 0.000 | 0.131 | 0.000 | 0.000 | | 0.000 | 0.000 | 0.000 | 0.000 | 0.000 | 0.000 | 0.000 |
| 6 | R007 | 1.000 | 0.004 | 0.000 | 1.000 | 0.000 | | 0.000 | 0.000 | 0.000 | 0.000 | 0.000 | 0.000 |
| 7 | LF3000 | 0.000 | 0.000 | 0.001 | 0.000 | 0.000 | 0.000 | | 0.999 | 0.000 | 0.990 | 0.255 | 0.041 |
| 8 | Turbofect | 0.000 | 0.000 | 0.000 | 0.000 | 0.000 | 0.000 | 0.999 | | 0.000 | 1.000 | 0.049 | 0.225 |
| 9 | Xfect | 0.000 | 0.000 | 0.000 | 0.000 | 0.000 | 0.000 | 0.000 | 0.000 | | 0.000 | 0.000 | 0.000 |
| 10 | JetOptimus | 0.000 | 0.000 | 0.000 | 0.000 | 0.000 | 0.000 | 0.990 | 1.000 | 0.000 | | 0.027 | 0.346 |
| 11 | TransIT X2 | 0.000 | 0.000 | 0.226 | 0.000 | 0.000 | 0.000 | 0.255 | 0.049 | 0.000 | 0.027 | | 0.000 |
| 12 | pDNA | 0.000 | 0.000 | 0.000 | 0.000 | 0.000 | 0.000 | 0.041 | 0.225 | 0.000 | 0.346 | 0.000 | |

### Supplementary Table 3

#### Statistical analysis of the transfection efficacy

Correlation matrixes between the numerical outputs of the various transfection assays, toxicity assays, and protein yields for both CHO and HEK293 cell lines. Pearson correlation are shown and significant correlations (p<.05) are highlighted in red.

#### Correlation matrixes: CHO

Correlations (Spreadsheet3 in 20220207 Transfection Correlation with means)
Marked correlations are significant at p < .05000
N=11 (Casewise deletion of missing data)

| Variable | Means | Std.Dev. | CHO adherent Luc | CHO suspension Luc | CHO SEAP suspension | CHO adherent GFP+ | CHO suspension GFP+ | CHO suspension GFP_low | CHO suspension GFP_med | CHO suspension GFP_high | CHO Protein Production | CHO Adherent Live | CHO Adherent Dead | CHO suspension Live | CHO suspension Dead | CHO adhesion MTS | CHO adhesion BrdU |
|---|---|---|---|---|---|---|---|---|---|---|---|---|---|---|---|---|---|
| CHO adherent Luc | 19928765 | 24852593 | 1.00 | -0.47 | -0.51 | 0.25 | -0.69 | -0.73 | -0.64 | -0.60 | -0.59 | -0.85 | 0.76 | 0.06 | -0.12 | -0.44 | -0.31 |
| CHO suspension Luc | 40235670 | 48744235 | -0.47 | 1.00 | 0.36 | 0.16 | 0.43 | 0.40 | 0.28 | 0.39 | 0.18 | 0.16 | -0.04 | 0.07 | -0.09 | 0.33 | -0.41 |
| CHO SEAP suspension | 22639 | 24510 | -0.51 | 0.36 | 1.00 | 0.33 | 0.85 | 0.69 | 0.80 | 0.93 | 0.88 | 0.53 | -0.40 | -0.04 | 0.06 | -0.22 | -0.06 |
| CHO adherent GFP+ | 71 | 14 | 0.25 | 0.16 | 0.33 | 1.00 | 0.21 | 0.21 | 0.30 | 0.23 | 0.34 | -0.31 | 0.17 | -0.20 | 0.08 | -0.34 | -0.01 |
| CHO suspension GFP+ | 34 | 26 | -0.69 | 0.43 | 0.85 | 0.21 | 1.00 | 0.95 | 0.97 | 0.97 | 0.86 | 0.60 | -0.46 | 0.10 | -0.09 | -0.19 | 0.16 |
| CHO suspension GFP_low | 13 | 9 | -0.73 | 0.40 | 0.69 | 0.21 | 0.95 | 1.00 | 0.96 | 0.87 | 0.78 | 0.57 | -0.45 | 0.11 | -0.12 | -0.11 | 0.31 |
| CHO suspension GFP_med | 12 | 11 | -0.64 | 0.28 | 0.80 | 0.30 | 0.97 | 0.96 | 1.00 | 0.94 | 0.89 | 0.58 | -0.50 | 0.09 | -0.09 | -0.23 | 0.35 |
| CHO suspension GFP_high | 9 | 8 | -0.60 | 0.39 | 0.93 | 0.23 | 0.97 | 0.87 | 0.94 | 1.00 | 0.86 | 0.55 | -0.42 | 0.12 | -0.11 | -0.25 | 0.10 |
| CHO Protein Production | 65 | 34 | -0.59 | 0.18 | 0.88 | 0.34 | 0.86 | 0.78 | 0.89 | 0.86 | 1.00 | 0.62 | -0.53 | -0.09 | 0.12 | -0.29 | 0.21 |
| CHO Adherent Live | 73 | 17 | -0.85 | 0.16 | 0.53 | -0.31 | 0.60 | 0.57 | 0.58 | 0.55 | 0.62 | 1.00 | -0.91 | -0.27 | 0.33 | 0.43 | 0.36 |
| CHO Adherent Dead | 6 | 5 | 0.76 | -0.04 | -0.40 | 0.17 | -0.46 | -0.45 | -0.50 | -0.42 | -0.53 | -0.91 | 1.00 | 0.41 | -0.45 | -0.51 | -0.55 |
| CHO suspension Live | 60 | 9 | 0.06 | 0.07 | -0.04 | -0.20 | 0.10 | 0.11 | 0.09 | 0.12 | -0.09 | -0.27 | 0.41 | 1.00 | -0.98 | -0.34 | 0.10 |
| CHO suspension Dead | 30 | 8 | -0.12 | -0.0[illegible] | 0.06 | 0.0[illegible] | -0.09 | -0.1[illegible] | -0.09 | -0.1[illegible] | 0.12 | 0.3[illegible] | -0.45 | -0.9[illegible] | 1.00 | 0.3[illegible] | -0.11 |
| CHO adhesion MTS | 5[illegible] | 26 | -0.44 | 0.33 | -0.2[illegible] | -0.34 | -0.1[illegible] | -0.11 | -0.2[illegible] | -0.25 | -0.2[illegible] | 0.43 | -0.5[illegible] | -0.34 | 0.3[illegible] | 1.00 | 0.14 |
| CHO adhesion BrdU | 1 | 0 | -0.31 | -0.41 | -0.06 | -0.0[illegible] | 0.16 | 0.31 | 0.35 | 0.10 | 0.21 | 0.36 | -0.55 | 0.1[illegible] | -0.1[illegible] | 0.14 | 1.00 |

### Correlation matrixes: HEK293

Correlations (Spreadsheet3 in 20220207 Transfection Correlation with means)
Marked correlations are significant at p < .05000
N=11 (Casewise deletion of missing data)

| Variable | Means | Std.Dev. | HEK adherent Luc | HEK suspension Luc | HEK SEAP suspension | HEK adherent GFP + | HEK suspension GFP+ | HEK suspension GFP_low | HEK suspension GFP_med | HEK suspension GFP_high | HEK Protein Production | HEK adherent Live | HEK adherent Dead | HEK suspension Live | HEK suspension Dead | HEK adhesion MTS | HEK adhesion BrdU |
|---|---|---|---|---|---|---|---|---|---|---|---|---|---|---|---|---|---|
| HEK adherent Luc | 43544619 | 28808114 | 1.00 | 0.48 | -0.84 | 0.41 | -0.48 | -0.20 | -0.53 | -0.42 | 0.06 | 0.19 | -0.25 | -0.17 | 0.11 | -0.48 | 0.01 |
| HEK suspension Luc | ######## | ######## | 0.48 | 1.00 | -0.46 | 0.34 | 0.26 | -0.44 | -0.08 | 0.41 | 0.47 | -0.05 | 0.12 | -0.01 | 0.00 | -0.38 | -0.50 |
| HEK SEAP suspension | 149568 | 112591 | -0.84 | -0.46 | 1.00 | -0.31 | 0.55 | 0.09 | 0.42 | 0.52 | -0.17 | 0.17 | -0.06 | 0.19 | -0.11 | 0.51 | -0.04 |
| HEK adherent GFP + | 63 | 19 | 0.41 | 0.34 | -0.31 | 1.00 | -0.07 | -0.18 | -0.42 | -0.09 | -0.39 | 0.05 | -0.11 | 0.18 | -0.17 | -0.09 | -0.51 |
| HEK suspension GFP + | 57 | 21 | -0.48 | 0.26 | 0.55 | -0.07 | 1.00 | -0.02 | 0.72 | 0.88 | 0.46 | -0.29 | 0.47 | -0.05 | 0.18 | 0.13 | -0.13 |
| HEK suspension GFP_low | 17 | 8 | -0.20 | -0.44 | 0.09 | -0.18 | -0.02 | 1.00 | 0.16 | -0.44 | 0.13 | -0.38 | 0.21 | -0.78 | 0.80 | -0.27 | 0.58 |
| HEK suspension GFP_med | 11 | 4 | -0.53 | -0.08 | 0.42 | -0.42 | 0.72 | 0.16 | 1.00 | 0.57 | 0.67 | -0.40 | 0.54 | -0.24 | 0.28 | 0.10 | 0.12 |
| HEK suspension GFP_high | 34 | 21 | -0.42 | 0.41 | 0.52 | -0.09 | 0.88 | -0.44 | 0.57 | 1.00 | 0.38 | -0.03 | 0.26 | 0.27 | -0.20 | 0.19 | -0.38 |
| HEK Protein Production | 64 | 16 | 0.06 | 0.47 | -0.17 | -0.39 | 0.46 | 0.13 | 0.67 | 0.38 | 1.00 | -0.36 | 0.44 | -0.54 | 0.49 | -0.44 | 0.11 |
| HEK adherent Live | 79 | 7 | 0.19 | -0.05 | 0.17 | 0.05 | -0.29 | -0.38 | -0.40 | -0.03 | -0.36 | 1.00 | -0.96 | 0.17 | -0.24 | 0.38 | -0.09 |
| HEK adherent Dead | 7 | 3 | -0.25 | 0.12 | -0.06 | -0.11 | 0.47 | 0.21 | 0.54 | 0.26 | 0.44 | -0.96 | 1.00 | -0.05 | 0.13 | -0.31 | 0.02 |
| HEK suspension Live | 62 | 12 | -0.17 | -0.01 | 0.19 | 0.18 | -0.05 | -0.78 | -0.24 | 0.27 | -0.54 | 0.17 | -0.05 | 1.00 | -0.97 | 0.39 | -0.55 |
| HEK suspension Dead | 23 | 11 | 0.11 | 0.00 | -0.11 | -0.17 | 0.18 | 0.80 | 0.28 | -0.20 | 0.49 | -0.24 | 0.13 | -0.97 | 1.00 | -0.28 | 0.59 |
| HEK adhesion MTS | 50 | 24 | -0.48 | -0.38 | 0.51 | -0.09 | 0.13 | -0.27 | 0.10 | 0.19 | -0.44 | 0.38 | -0.31 | 0.39 | -0.28 | 1.00 | -0.04 |
| HEK adhesion BrdU | 1 | 0 | 0.01 | -0.50 | -0.04 | -0.51 | -0.13 | 0.58 | 0.12 | -0.38 | 0.11 | -0.09 | 0.02 | -0.55 | 0.59 | -0.04 | 1.00 |
